# A natural mechanism of eukaryotic horizontal gene transfer

**DOI:** 10.1101/2025.02.28.640899

**Authors:** Andrew S. Urquhart, Samuel O’Donnell, Emile Gluck-Thaler, Aaron A. Vogan

## Abstract

Horizontal gene transfer (HGT) is an important and potentially frequent process impacting the evolutionary trajectory of eukaryotic species. Yet to date, the mechanism of HGT among eukaryotes has remained a mystery. Here, we demonstrate that *Starships*, a newly discovered group of cargo-mobilizing transposable elements, are active vectors of horizontally transferred DNA among eukaryotic species. We find that multiple *Starships* simultaneously transfer themselves and their cargo within and between fungal genera separated by upwards of 100 million years of evolution. Transferred Starships are recovered in strains with and without selection for particular *Starship*-encoded phenotypes. Using whole genome sequencing, we exclude alternative hypotheses such as heterokaryosis, parasex or the random uptake of exogenous DNA by showing that only *Starship* DNA is precisely transferred. Furthermore, we demonstrate that the introduction of the *Hephaestus Starship* into a new species increases metal resistance due to the genes carried by this element. Between 18-27% of all *Starships* in our focal genus, *Paecilomyces,* have evidence of horizontal transfer under natural conditions, including to other fungal genera, revealing *Starship*-mediated HGT to be a widespread and frequent phenomenon across filamentous fungi. This study identifies the first active genetic mechanism for HGT between eukaryotic species.

## Introduction

HGT is a consequential means of genetic exchange predicted to occur between eukaryotes, but the mystery of how it happens across this domain of life has remained unsolved for decades. Unlike vertical inheritance, HGT involves the transfer of DNA between individuals in the absence of sex. Advances in sequencing coupled with comparative genomics have revealed many strong cases of HGT across a wide range of eukaryotes [1–3]. For example, an antifreeze-encoding protein gene was horizontally transferred between two fish [4], while a gene required to detoxify plant defence compounds was obtained by a whitefly from plants [5–7]. Yet while we often conclude *that* HGT has happened, and sometimes identify plausible adaptive explanations for *why* it may have happened, we almost never know *how* it happened [8]. Thus, we have limited insight into how the movement of genetic material across eukaryotic species is achieved and few opportunities to study the biology of eukaryotic HGT under laboratory settings.

Fungi are ideal model systems for investigating eukaryotic HGT due to their tractability and the impetus to study them for their societal and environmental importance. Because fungal interactions with both plants and animals pose threats and opportunities for food security and public health, understanding how fungi adapt to new hosts and environments is an ecological and economic priority [9]. One major reason to be concerned about fungal HGT is that it provides major adaptive gains (e.g. acquisition of a new virulence factor) with a single mutational event [10]. This has led to the emergence of new pathogens of both plants and animals like *Metarhizium robertsii,* an insect pathogen whose shift to pathogenicity was facilitated by HGT [11]. The first identified examples of putative fungal HGTs were likely acquisitions of bacterial genes whose evolutionary histories clearly differ from the surrounding genome [12]. More recently, a growing number of genes transferred between fungi have been identified through comparative genomics and phylogenetics [13,14]. Notable examples include the *toxA* virulence gene whose transfer resulted in the emergence of a new wheat pathogen [15], the transfer of multiple *Fusarium* virulence factors [16], and the transfer of the sterigmatocystin biosynthetic gene cluster which resulted in the *de novo* gain of an entire secondary metabolite pathway [17]. However, as with other eukaryotes, interspecies DNA transfers among fungi lack a mechanism, precluding a basic understanding of HGT biology.

One striking feature about fungal HGT is that not only is there phylogenetic evidence of individual genes having transferred, but entire genomic regions spanning hundreds of kilobases [18]. One untested hypothesis to explain these data is that transposable elements carrying large amounts of sequence as cargo are vectors of eukaryote-eukaryote HGT [19]. This hypothesis is supported by the striking distribution of near identical massive transposons across different fungal species [20–22]. These transposons, which we have named *Starships,* are a recently discovered group of transposable elements found in filamentous Ascomycete fungi (*Pezizomycotina*), a large group of ecologically diverse fungi consisting of more than 82,000 described species [21–23]. *Starships* are mobilised by a tyrosine recombinase, the “captain”, which is always found as the first gene in the element [21]. *Starships* possess an unprecedented capacity among eukaryotic DNA elements to carry additional genetic cargo that encode organismal phenotypes beyond transposition-related functions [24]. There are a growing number of *Starships* in taxonomically diverse species spanning the *Pezizomycotina* for which genomic evidence indicates past movement through horizontal transfer [21,24–28]. As one example, a set of *Starships* carrying a cluster of genes providing resistance to formaldehyde appear to have been transferred at least nine times among different fungal species [29].

While mechanisms of DNA exchange are well established in the bacterial domain, a decisive explanation for how DNA moves horizontally among eukaryotes is missing [8,30,31]. Here, we show that *Starship* transposons transfer between species from the same and different genera under simple co-culture conditions. We rule out alternative explanations of heterokaryosis, introgression, random DNA uptake, and parasexuality through a combination of selective culturing methods and whole genome sequencing of recipients. We leveraged a large genomic dataset and found that at minimum 18-27% of all *Starships* in our genus of focus, *Paecilomyces*, have undergone at least one inter-species transfer, underscoring the prevalence and incidence of *Starship-*mediated HGT in natural environments. Thus, we demonstrate that *Starships* are a mode of genetic exchange in fungi, and provide the first active mechanism of inter-eukaryotic HGT.

## Results

### The *Starship* transposon *Hephaestus* is horizontally transferred within and between species

One of the strongest cases for eukaryote-eukaryote HGT is the repeated transfer of the *Hephaestus Starship* [25]. *Hephaestus* is a large transposon belonging to the *Starship* superfamily that carries resistance to multiple metal ions, first identified in the environmental fungus *Paecilomyces variotii* [25]. Near-identical copies of *Hephaestus* are present in two other *Paecilomyces* species, *P. paravariotii* [32] and *P. lecythidis* [21], and in another genus, *Penicillium chermesium* [29]. However, in the absence of experimental evidence, alternative explanations remain for why identical copies of this transposon are found in different species. For example, it has been suggested that apparent *Starship*-mediated HGTs might result from hybridisation followed by the subsequent loss of non-*Starship* DNA [33]. Given that *Hephaestus* is predicted to have undergone multiple horizontal gene transfers, it represents the ideal candidate for testing the hypothesis that *Starships* are a previously hidden mechanism of eukaryotic HGT.

We tested whether the horizontal transfer of *Hephaestus* occurs under lab conditions (Figure 1A). First, we tagged *Hephaestus* with a hygromycin resistance gene in a leucine auxotroph background of *P. variotii* CBS 144490 [34]. This resulted in a strain, hereafter the “donor strain”, resistant to hygromycin but auxotrophic for leucine i.e. unable to grow in the absence of supplemental leucine. Thus, when paired to a wild type “acceptor strain” sensitive to hygromycin but prototrophic for leucine, the predicted result of a successful *Starship* transfer would be a hygromycin-resistant and leucine-prototrophic “recipient strain”. We co-cultured the donor strain with three different acceptor strains of increasing phylogenetic distance. The three acceptor strains used were wild type isogenic *P. variotii* CBS 144490 (same species), *P. paravariotii* FRR 5287 (different species, which has a previously inferred horizontal transfer of the *Hephaestus Starship* [32]), and a strain belonging to a different genus *Aspergillus fumigatus* A1160. All acceptor strains are leucine-prototrophic and sensitive to hygromycin (Figure 1B). We obtained multiple hygromycin-resistant recipient strains for each strain pairing (Figure 1B). In total 138 pairings were made, and of these 10/16 (62.5%) of the *P. variotii* CBS 144490, 95/98 (96.9%) of the *P. paravariotii* FRR 5287 and 4/24 (16.7%) of the *A. fumigatus* A1160 pairings resulted in recipient colonies with the predicted phenotypes indicative of *Starship*-mediated horizontal transfer (Table S1).

**Figure 1:**
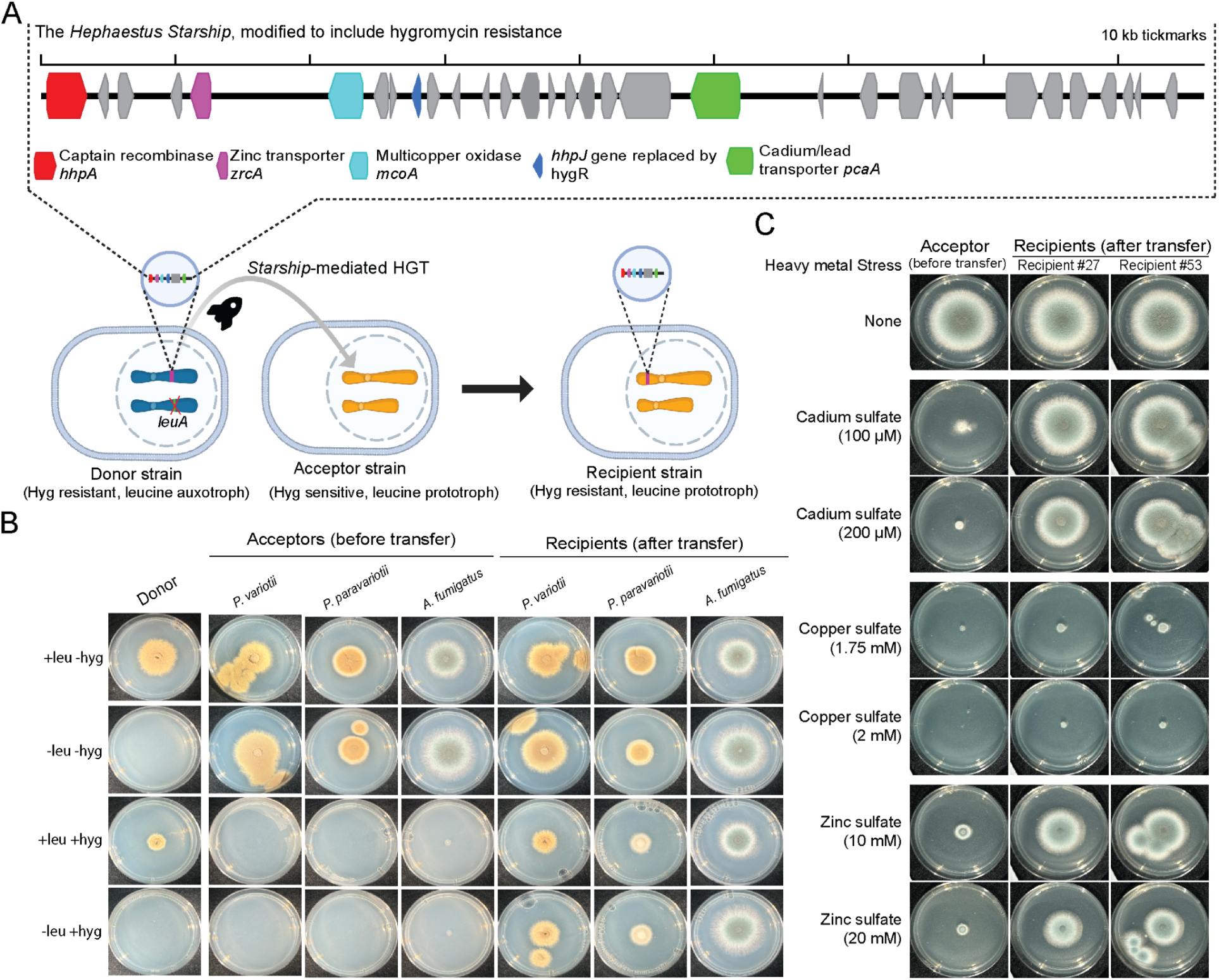
Horizontal transfer of the *Hephaestus Starship* across species and genera. **A)** An experimental screen for *Starship*-mediated HGT reveals the simultaneous transfer of both engineered and naturally occuring adaptive phenotypes. A genetically-modified “donor” strain was constructed to carry a hygromycin antibiotic resistance gene (hygR) on the *Hephaestus Starship* and additionally to be unable to grow in the absence of the amino acid leucine (auxotroph). Horizontal gene transfer of the *Starship* enables the recipient to grow in the presence of hygromycin. In addition to carrying the introduced antibiotic resistance *Hephaestus* naturally contains a number of genes involved in metal tolerance. Abbreviations Hyg = hygromycin, leu = leucine, *LeuA* KO *=* deletion of the *leuA* gene required for leucine biosynthesis. **B)** Gain of antibiotic resistance following acquisition of hygR-tagged *Hephaestus*. Growth of donor, acceptor and recipient strains on minimal media with and without leucine and hygromycin. **C)** Gain of Cu^2+^, Cd^2+^ and Zn^2+^ resistance by *A. fumigatus* following acquisition of hygR-tagged *Hephaestus*.

### The inter-genus transfer of *Hephaestus* from *Paecilomyces* to *Aspergillus* increases metal resistance

One of the features that make *Starships* distinct from other eukaryotic transposons is that they transfer large segments of protein-coding DNA in a single event (typically dozens of genes spanning as much as 700 kb, but more commonly ∼100 kb [35]), which provides an opportunity for recipients to simultaneously acquire multiple adaptive phenotypes [24]. *Hephaestus* provides resistance to at least five different metal ions, raising the question of whether these different phenotypes would still be expressed after a *bona fide* transfer to a species in a different genus [25]. Of the three acceptors we used, *P. variotii* CBS144490 and *P. paravariotii* FRR 5287 already have a native copy of *Hephaestus* whereas *A. fumigatus* A1160 does not. As such, the effect of acquiring *Hephaestus* on metal resistance was assessed in the *A. fumigatus* recipients. Indeed, we found that the *de novo* acquisition of *Hephaestus* by *A. fumigatus* increases resistance to the three metal ions that we tested (Cd^2+^, Zn^2+^, Cu^2+^; Figure 1C). The expression of multiple metal resistance phenotypes with native promoters and an engineered antibiotic resistance phenotype in recipient strains supports the conclusion that *Starships* have capacity to make complex contributions to fungal phenotypes and are drivers of multivariate adaptation across fungal genera.

### Whole genome sequencing confirms the stable integration of *Hephaestus* post-transfer

The increased hygromycin resistance phenotype of the recipient colonies suggests these strains now carry the hygR-tagged *Hephaestus Starship* acquired from the donor. To confirm that hygromycin resistance was indeed conferred by the horizontal transfer of *Hephaestus*, we used whole genome Illumina shotgun sequencing of the recipients in our co-cultivation experiments. Both *P. variotii* CBS 144490 and *P. paravariotii* FRR 5287 have a native-copy of *Hephaestus* so the expected result of a *Starship*-mediated HGT into these strains is for *Hephaestus* copy number to increase from 1 to 2 and for this second copy to be located at a different position in the genome compared to both the donor and recipient. In most cases we saw an approximate doubling of the read depth across *Hephaestus* relative to the genome as anticipated (Figure 2A). However, in two cases where *P. variotii* CBS 144490 was the acceptor strain, *Hephaestus* appeared in 3 copies in the recipient indicating that 2 copies of *Hephaestus* had been transferred and subsequently integrated into the genome (Figure 2A, Figure S1; Table S2). Similarly, with an *A. fumigatus* A1160 acceptor (which possesses no native copies of *Hephaestus*), *Hephaestus* went from 0 copies to either 1 to 2 copies (Figure 2A, Figure S1; Table S2). *De novo* genome assembly of the two *A. fumigatus* recipients that had obtained the *Starship* indicated that *Hephaestus* is newly embedded within their genomes. Thus unambiguously connecting the antibiotic and heavy metal resistance phenotypes with the transfer of *Hephaestus* (Figure 2B, Figure S2). As expected, with few exceptions, native copies of *Hephaestus* were maintained at their original locus while transferred copies integrated into new insertion sites with the canonical TTAC(N7)A motif (Figure 2C and D, Figure S3, see supplemental results “***Hephaestus* is integrated into the genome following HGT”**).

**Figure 2:**
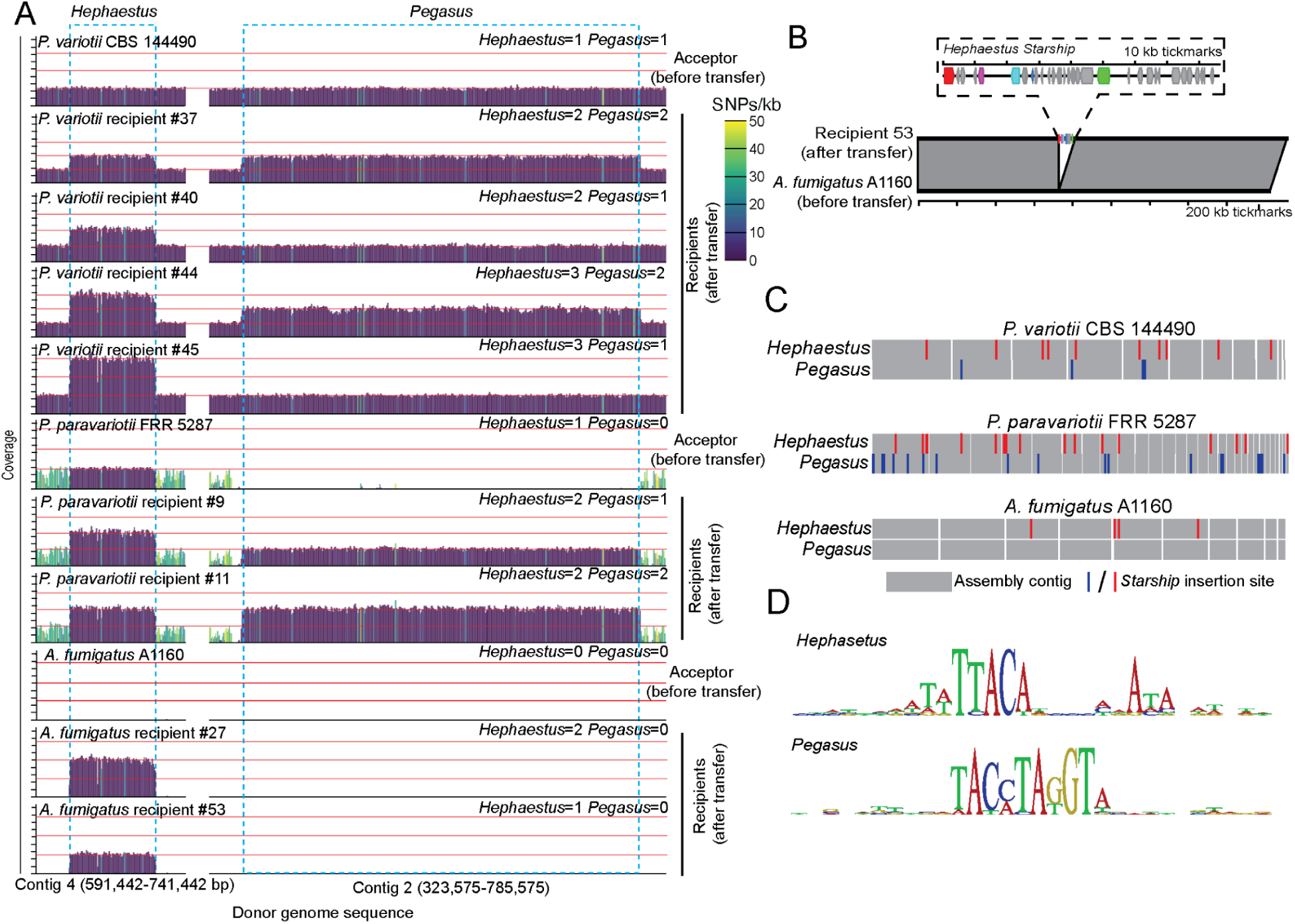
Genome sequencing reveals the co-HGT of two *Starships*, *Hephaestus* and *Pegasus*. **A)** Read-mapping of Illumina sequencing reads derived from recipient strains to the *Starships Hephaestus* and *Pegasus* and flanking DNA in the donor genome (ie. *P. variotii* CBS 144490). Y axis indicates read coverage (from 0 to 4500 reads mapped per 1 kb window). Colour indicates the number of SNPs within each 1 kb window. Red horizontal lines indicate estimates of copy number from 1 to 3. **B)** Comparison between the *A. fumigatus* recipient #53 contig containing the integrated *Hephaestus* and the wild type *A. fumigatus* A1160 chromosome 4 (nucleotide position 1495672 to 3322498). **C)** Position of *Hephaestus* and *Pegasus* insertions within the recipient genomes. Grey rectangles represent sequencing contigs. For *P. paravariotii* FRR 5287 the largest 29 contigs representing 98.5% of the total assembly are shown. **D)** Sequence logo of target sites at which *Hephaestus* and *Pegasus* were found within the recipient strains.

### *Starship Pegasus* co-transfers with *Hephaestus*

In interpreting the results of our HGT transfer experiments, we explicitly considered alternative explanations (beyond simple spontaneous resistance) for the emergence of apparent recipient phenotypes, i.e. strains that are hygromycin-resistant and able to grow without leucine. These alternative scenarios, including introgression, parasexual recombination, exogenous DNA uptake and hybridization, would result in additional DNA beyond the *Starship* being transferred (Figure 2A, Figure S1). Thus, to rule out these alternative hypotheses we mapped the Illumina reads from the recipient strains back to the donor genome to demonstrate that they precisely contain *Starship* DNA embedded in their genome and no other donor DNA (**see supplementary discussion “extended arguments for why HGT is the most likely explanation”**).

Outside of *Hephaestus*, a single additional region with signatures of HGT is observed in some recipient strains. However it does not correspond to random genomic sequence (as would be expected if transfer was achieved through the random uptake of exogenous DNA), but rather to a second *Starship* which we had previously identified within the *P. variotii* CBS 144490 genome and named *Pegasus* [21]. Neither *P. paravariotii* FRR 5287 nor *A. fumigatus* naturally contain *Pegasus.* The movement of *Pegasus* is remarkable given the lack of selection for its maintenance in recipient strains (unlike hygR-tagged *Hephaestus*). These data support a prior hypothesis based on bioinformatic analyses that *Hephaestus* and *Pegasus* horizontally transfer together, given their apparent simultaneous appearance in the genomes of wild-type environmental strains from different species [21]. We note that both *Hephaestus* and *Pegasus* are the only two *Starships* found in the donor strain *P. variotii* CBS 144490. *Pegasus* is the largest of all *Starships* detected across the *Paecilomyces* genus, totalling 403.7 kb in length (see below).

The transfer of *Hephaestus* does not appear to depend on the transfer of *Pegasus.* Of the 26 recipient strains that were sequenced with Illumina: 50% (4/8) derived from *P. variotii* CBS 144490 obtained 1 additional copy of *Pegasus*, and 100% (15/15) of recipient strains derived from *P. paravariotii* FRR 5287 obtained at least 1 copy of *Pegasus* with 26.7% (4/15) obtaining 2 copies (Figure 2A, Figure S1, Table S2). We did not observe the co-transfer of *Pegasus* into *A. fumigatus*, but fewer of these recipients were sequenced (Figure 2A, Figure S1, Table S2). As with *Hephaestus*, we determined integration sites in the recipient genomes by mapping reads to the edges of *Pegasus*. These target sites were consistent with the 10 bp conserved target sites observed for a mini-*Pegasus Starship* (Figure 2D, Figure S4, **see supplemental results “characterisation of the *Pegasus Starship”***). In some cases *Hephaestus* is found nested within *Pegasus*. It is impossible to know *post hoc* if this was a nested transfer (i.e. *Hephaestus* nested prior to transfer) or a sequential transfer with *Hephaestus* coincidentally finding a target site within *Pegasus.* Since *Pegasus* does not have a selectable marker, we do not know whether its transfer is dependent on *Hephaestus*.

The finding that two *Starships* have horizontally transferred is intriguing given the known complex interactions between bacterial mobile elements that lead to both conflict and cooperation between different elements [36]. This includes the existence of “Hitcher Genetic Elements” which require the help of other elements to transfer [37]. Given our observation of consistent co-transfers, future studies may uncover similar dynamics between *Starship* elements as well as the specific mechanism of *Starship* movement between strains.

### *Starship*-mediated HGT is a prevalent natural phenomenon

The prevalence and incidence of eukaryotic HGT is strongly debated [8]. To provide greater ecological and evolutionary context to the experimental demonstration of HGT under laboratory conditions, we estimated the extent of naturally-occurring *Starship*-mediated HGT across the *Paecilomyces* genus using comparative genomics. We conducted automated and manual annotations of *Starships* across *Paecilomyces* to find that this genus hosts a diverse repertoire of 33 distinct *Starships*, ranging from 16-403 kb in size (71.5 kb on average) and carrying between 1-123 protein-coding genes (18 genes on average; **see supplemental results “Survey of *Starships* in *Paecilomyces”***; Table S3; Figure S6-S18. We systematically searched for recently transferred copies of these elements across the *Pezizomycotina*, the largest subphylum of filamentous fungi predicted to be the primary hosts of *Starships* (Table S3.5) [35]. We found that upwards of 9/33 *Starships* (27.3%; ranging from 52.0 kb - 403 kb) have evidence of horizontal transfer either to or from various *Paecillomyces* species, with 6 (18.2%) having crossed genus-level boundaries (Figure 3; Table S3.6). We consider some of the within genus transfers to only be likely cases of HGT, since it can be challenging to completely rule out alternative explanations such as introgression or incomplete lineage sorting among more closely related species. However, these alternative explanations lose power when evaluating other cases of within genus transfers, such as *Pegasus*, where the observed sequence similarity among transferred *Starships* vastly exceeds background levels across the entire genome. Among the 9 *Starships* with evidence of transfer, 4 were previously identified (*Hephaestus* [25] (Figure S7), *Chrysaor* [29] (Figure S8), *Prometheus* [29] (Figure S9), and *Pegasus* [21] (Figure S10)) and most encode conditionally beneficial cargo such as genes conferring heavy metal resistance and formaldehyde detoxification. The 5 additional *Starships* with evidence of horizontal transfer are described here for the first time, thereby roughly doubling previous assessments of horizontally-transferable *Starships* within the *Paecilomyces* genus: *Demeter* (Figure S11), *Hades* (Figure S12), *Hera* (Figure S13), *Hestia* (Figure S14) and *Keraunos* (Figure S15).

**Figure 3:**
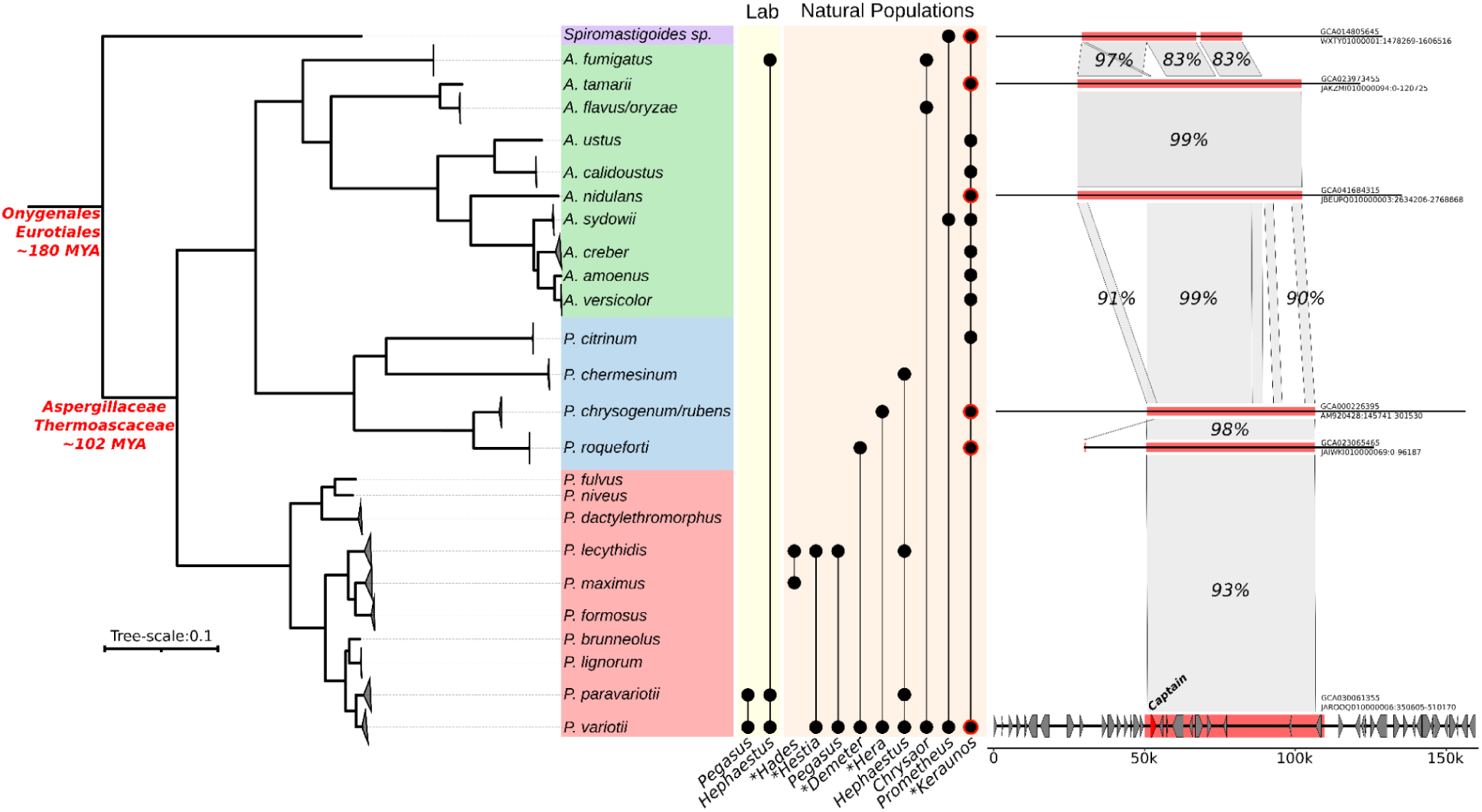
*Starships* transfer across fungal species and genera. The maximum likelihood phylogeny (left) was generated using conserved BUSCO orthologs from genomes containing predicted *Starships* detected in the horizontal gene transfer (HGT) screen. All branches have 100% ultra-fast bootstrap and aLRT support. Estimated branching times in red are derived using TimeTree [40]. Each black dot in the plot to the immediate right of the phylogeny represents the presence of a *Paecilomyces*-derived *Starship* in at least one genome from the associated species in the phylogeny as detected using BLAST. *Starships* in the “Lab” columns summarize the findings of our experimental transfer data, while *Starships* in the “Natural populations” columns summarize the findings of our comparative genomic screen across the *Pezizomycotina* (n=10,010 genomes). * indicates newly described *Starships. T*he synteny alignments (right) summarize nucleotide identity among copies of the *Keraunos Starship* with evidence of horizontal transfer in species for which we were able to recover complete and unfragmented copies of this element. Predicted gene models are symbolized as gray arrows (no *de novo* gene predictions were performed for assemblies outside of *Paecilomyces*).

We note that our estimates of *Starship* diversity within *Paecilomyces* and thus our subsequent estimates of horizontal transfer rates are conservative, as the vast majority of *Starships* within a given species typically segregate at low frequencies [38]. *Starships* are furthermore more likely to be annotated if using a dataset containing multiple contiguous long-read assemblies from the same species [38]. We therefore do not expect to have recovered the complete set of *Starships* from *Paecilomyces*, since our dataset mostly consists of short-read assemblies with, in many cases, few sequenced strains per species. For example, we found that, on average, only 19% of all annotated captains in each genome were associated with a full-length *Starship* element (Figure S18). Additionally, depending on the annotation method used (either BLAST-based searches of known elements or *de novo* starfish predictions), between 15-24 of the 33 *Paecilomyces Starships* are only ever found in a single strain, as are the vast majority of *Starships* with evidence of horizontal transfer in other *Pezizomycotina* species. We note this bias would not impact our main conclusions, other than potentially leading us to underestimate the prevalence of *Starship* horizontal transfer. Despite these biases, most of the transferred *Starships* are restricted to the *Eurotiales* (the taxonomic order containing our query genus), supporting previous findings that the frequency of *Starship*-mediated HGT likely decreases with increasing phylogenetic distance [39]. Yet even so, several transfers involving near identical elements have occurred between distantly related species, including between the orders *Eurotiales* and *Onygenales*. The long distance transfers suggest that *Starships* are able to mediate horizontal transfer across species separated by upwards of 180 million years of evolution (Figure 3), roughly equivalent to a transfer event between monotremes and eutherian mammals [40].

Our estimate that between 18.2-27.3% of *Paecilomyces Starships* show evidence of recent HGT in natural fungal populations (including the HGT of relatively rare *Starships* in our dataset) suggests *Starship*-mediated HGT is a commonly occurring natural phenomenon, as opposed to either an experimentally-induced artifact or a rare event. These observations align with the fact that *Starship*-mediated HGT in the lab utilized near-natural conditions: strains were merely co-cultured on standard fungal growth media, as opposed to imposing highly artificial interventions such as harsh chemical treatment and/or the removal of the cell wall, as is typically done during conventional fungal transformations. Furthermore, in the case of *Pegasus,* the *Starship* transfer and subsequent maintenance occurred without directed selection for any particular cargo carried by this element, highlighting the fact that *Starship* transfer, like any other mutational event, occurs prior to selection. Although we focused on specific taxa, *Starship*-mediated horizontal transfer is not limited to the *Eurotiales* or *Paecilomyces*: indeed, reports of *Starships* with evidence of horizontal transfer appear in multiple species from taxonomic classes across the *Pezizomycotina* [26,27,41]. We thus conclude that *Starships* must now be included in the repertoire of naturally-occurring and experimentally demonstrated mechanisms of fungal gene exchange (Figure 3B). This places *Starship* transfer alongside processes such as sexual recombination first described in the 1820’s [42,43], parasex first shown experimentally in the 1950’s [44] (**see supplementary discussion “Horizontal Chromosome Transfer: a special case of parasex?”**), and natural transformation first shown experimentally in the 1970’s [45].

**Figure 4:**
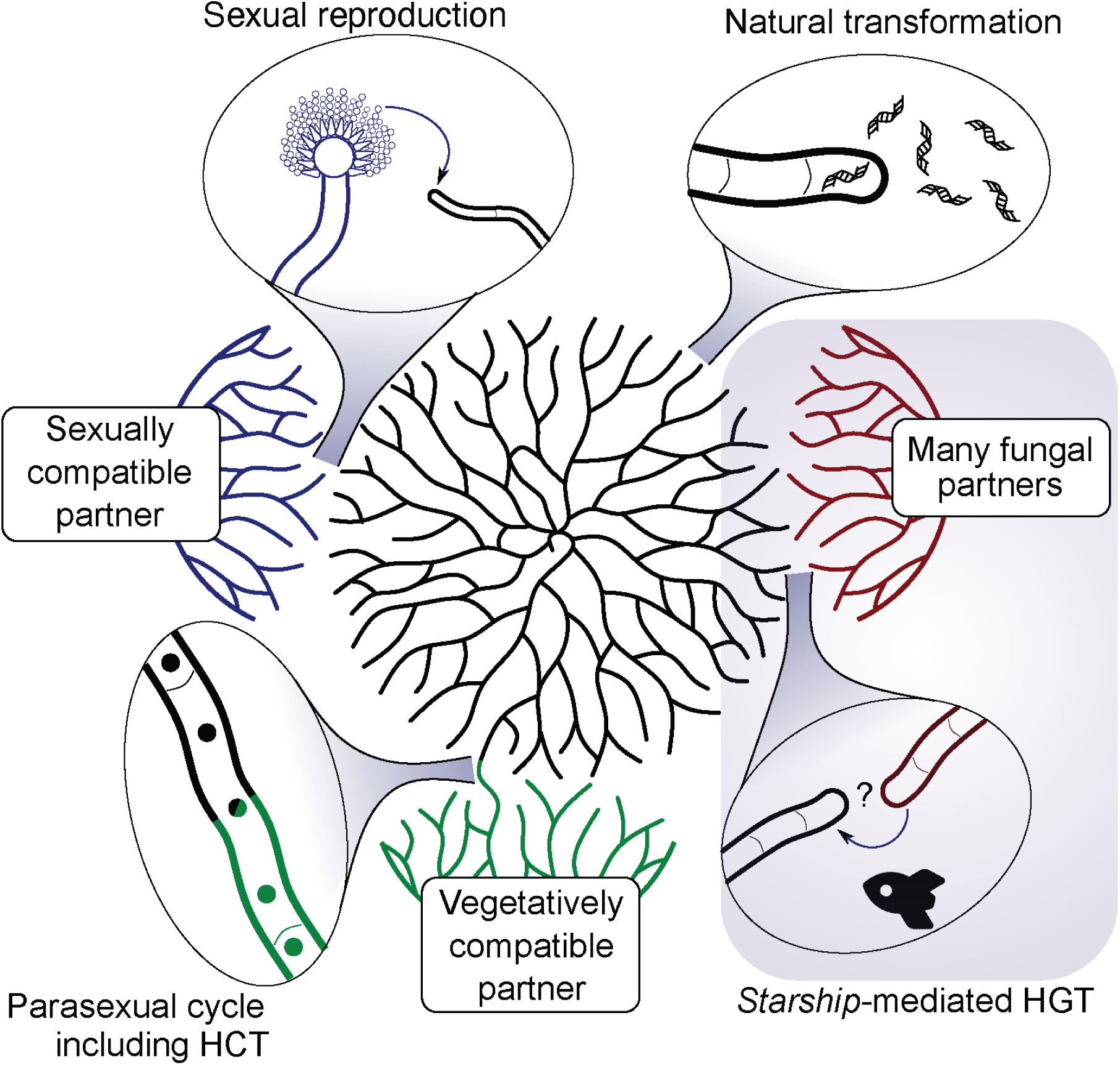
Mechanisms of genetic exchange in fungi. *Starship*-mediated horizontal gene transfer (HGT) adds a new mechanism to the repertoire of established and naturally-occurring processes of genetic exchange in fungi. HCT = horizontal chromosome transfer.

## Conclusion

We demonstrate that *Starship* giant transposable elements are a mechanism of horizontal transfer in eukaryotes. This provides an answer to what has been a decades-old mystery: how DNA is transferred between fungal species [46]. Giant transposable element-mediated HGT might be relevant to eukaryote evolution more broadly. *Mavericks* are giant virus-like transposons first discovered in the mid 2000’s within the genomes of a wide diversity of eukaryotes [47,48]. Recently, it has been shown that *Maverick* transposons have been horizontally transferred between nematode species and furthermore that *Mavericks* carry a putative cargo gene which is not required for mobility [20]. Future work may reveal that *Mavericks* or some other currently unknown transposons are analogs of *Starships* in animals and plants. The experiments presented here place the study of eukaryotic HGT at a ‘coming of age’ moment and prepare us to more adequately address fungal threats and opportunities head on with mechanistic knowledge of how the exchange of genetic information is achieved across species boundaries.

## Methods

### Fungal strains and media

*Paecilomyces variotii* strains CBS 144490 (*Hephaestus+*, *Pegasus*+), *Paecilomyces paravariotii* strain FRR 5287 (*Hephaestus+*, *Pegasus*-) and *Aspergillus fumigatus* A1160 (*Hephaestus-*, *Pegasus*-) [49,50] were cultured on potato dextrose agar (PDA) supplemented with 380 mg/L leucine except for where selection against leucine auxotrophs was required in which case minimal medium was used [51] or where antibiotic selection was required in which case the medium was supplemented with either hygromycin (100 µg/ml for *Paecilomyces* and 250 µg/ml *Aspergillus*) or G418 (200 µg/ml). For confirming hygromycin resistance and leucine auxotrophy phenotypes *Paecilomyces* was incubated for 5 days and *Aspergillus* for 3 days both at 30°C.

### DNA extraction

DNA was extracted from lyophilized mycelium grown overnight at 30°C in PDA following a modified version of that followed by Pitken *et al.* [52]. Samples were homogenized by bead beating, suspended in 1 ml of CTAB buffer (100 mM Tris-HCl pH 7.5, 0.7 M NaCl, 10 mM EDTA pH 8, 1% CTAB, 1% β-mercaptoethanol) then incubated at 65°C for 30 minutes. 500 µl was then added to an equal volume of chloroform and centrifuged at maximum speed in a benchtop centrifuge for 10 min. The aqueous phase was mixed with an equal volume of isopropanol to precipitate the DNA which was subsequently pelleted, washed with 70% ethanol and air dried before being resuspended in H_2_O.

### Nanopore sequencing and assembly of *P. variotii* CBS 144490

In the previous *P. variotii* CBS 144490 genome assembly based on Illumina reads [34] the *Pegasus Starship* was highly fragmented [21]. Thus we resequenced this strain using Oxford Nanopore at the Central Analytical Research Facility at the Queensland University of Technology. DNA was extracted from two runs, one using unsheared DNA, the LSK110 ligation sequencing kit, and Minion flow cell chemistry R9.4.1. The second was conducted similarly but using DNA sheared to approximately 20 kb using a 26G needle. Assembly and polishing steps were conducted in Galaxy [53]. The reads were assembled using Flye (Galaxy version 2.9 [54]) with two polishing iterations and a minimum overlap of 7,044 bp between reads. Previously generated Illumina reads (SRA SRX4939669) were mapped to the genome using Bowtie 2 (Galaxy version 2.4.5 [55]) to correct errors associated with Nanopore sequencing [56] using Pilon [57]. Gene annotations were transferred from the JGI assembly using liftoff version v1.5.1 [58].

### Cloning

#### 1. Construction of mini-*Pegasus*

A minimal version of *Pegasus* was constructed analogous to a minimal *Hephaestus* construct which we reported previously (Figure S4) [21]. Minimal *Pegasus* consisted of 4,770 bp of 5’ terminal sequence including the captain sequence, the *zrcA* gene used as a selectable marker and 2060 bp of 3’ terminal sequence. The minimal *Starship* was nested inside the hygromycin selectable marker. To achieve this mini-*Pegasus* was cloned within an intron (the second intron of the gene model encoding protein JGI ID 278152 from CBS 144490 [34]) added to the HYG resistance cassette which naturally contains the *Pegasus* direct repeat sequence previously observed [21].

The first half of the hygromycin resistance cassette was amplified from plasmid PMAI6 [59] with primers AP244+AP584 (Primer sequences are given in Table S4). The 5’ of *Pegasus* was amplified in two halves with primer pairs AP595+AP593 (subsequently amplified with AP596+AP593) and AP586+AP592. The *zrcA* selectable marker that confers resistance to Zn^2+^ ions was amplified with AP587+AP588, the 3’ of *Pegasus* was amplified with AP589+AP590 and the second half of the hygromycin resistance cassette was amplified with AP591+AP243. All fragments were cloned into the plasmid PLAUB36 [21] digested with EcoRI and HindIII, via homologous recombination in yeast. The required intron sequence was included within the primer sequences. The design of the construct was imperfect as it resulted in an inadvertent duplication of the 5’ direct repeat, and as a result leaves a copy of the direct repeat inside the intron following excision. However, this additional copy of the direct repeat did not impact the function of the intron (as demonstrated through hygromycin resistance) so redesigning the construct was unnecessary.

#### 2. Construction of a *leuA* gene replacement (KO) strain

A construct was made to delete the *leuA* homolog of *Paecilomyces* CBS 144490. Loss of this gene results in leucine auxotrophy in *Paecilomyces variotii* as in other fungi [34]. 1681 bp 5’ to the gene was amplified with primers ASW7+ASW8, the G418 resistance cassette of PMAI2 [59] was amplified with ASW9+ASW10 and 1655 bp 3’ to the gene was amplified with primers ASW11+ASW12. The fragments were combined into plasmid PLAUB36 [21] digested with EcoRI and HindIII via homologous recombination in yeast.

#### 3. Construction of *hhpJ* KO

In order to tag the *Hephaestus Starship* we targeted a hygromycin resistance cassette by homologous recombination into the *hhpJ* locus in *Hephaestus*. 2,307 bp of sequence 5’ to the gene were amplified with primers ASW1+ASW2, the HYGR resistance cassette of PMAI6 was amplified with ASW3+ASW4 and 2,233 bp 3’ to the gene were amplified with primers ASW5+ASW6. The fragments were combined into plasmid PLAUB36 [21] digested with EcoRI and HindIII via homologous recombination in yeast.

### Transformation of *Paecilomyces variotii* and screening for homologous integration

*Paecilomyces variotii* was transformed using *Agrobacterium* strain EHA105 [34]. Integration in the *leuA* locus was screening for phenotypically via plating transformants on to media with and without leucine. Integration in the *hhpJ* locus within *Hephaestus* was assayed via PCR with primers pair ASW73+ASW74 which amplify a fragment internal to the *hhpJ* gene. The genotype of the donor strain was confirmed using Illumina sequencing and mapping the reads to the *leuA* and *hhpJ* regions using Bowtie 2 [55] to confirm their absence (Figure S5).

### Transposition of *Pegasus* within genome

Single germlings of the mini-*Pegasus* transformants were placed at the center of a 90 mm PDA plate and allowed to grow for two weeks at 30°C. At that time the spores were harvested from the plate in 5 ml of H_2_O and 1 ml aliquots of spore suspension were replated onto PDA plates containing hygromycin. Emergent hygromycin resistant colonies were picked and then Illumina sequenced as a pool.

### Coculture of donor and acceptor strains to produce recipient strains

To demonstrate the horizontal movement of *Starships* between strains we first generated a “Donor strain”. This strain was transformed with the *hhpJ* (hygromycin selection) and *leuA* (G418 selection) knockout constructs. This strain is unable to grow in the absence of leucine. We then paired this strain with various wild-type acceptor strains including the isogenic wild type, *P. paravariotii,* and *Aspergillus fumigatus* A1160. Pairings were made by streaking both strains on top of each other in a straight line down the center of a 90 mm PDA petri dish then leaving the strains to grow together at 20°C for four weeks. After which time 1/5th of the spores from the plate were transferred onto media lacking leucine with hygromycin and incubated until recipients emerged. Plates were scored in a binary manner as either having or not having recipient growth. For each plate a representative transformant was selected and purified by streaking spores onto a petri dish and excising a single germling after overnight incubation. The genotype of the recipients was validated either by Illumina genome sequencing or for some cases involving FRR 5287 by PCR. For PCR a region 801 bp upstream of the 5’ TTAC direct repeat flanking *Hephaestus* in *P. variotii* CBS 144490 was amplified with primer pair ASW269+ASW270. The amplified region was polymorphic between FRR 5287 with an EcoRI site unique to *P. variotii* CBS 144490. As such PCRs could be digested with EcoRI to determine the background genotype of the recipient. The presence of the hygromycin and *leuA* alleles were confirmed using a multiplex PCR with primers ASW160+ASW161 amplifying a fragment of the hygromycin resistance marker and ASW66+ASW67 amplifying a fragment of the *leuA* gene.

### Illumina sequencing analysis

DNA was sequenced via 150 bp paired-end libraries prepared using either an Illumina DNA prep (M) kit at the Australian Genome Research Facility (for mini-*Pegasus*), a Watchmaker PCR-free kit at Macrogen (most recipient strains) or at Novogene (the donor strain and recipient strains 1, 2, 7 and 8).

Reads of each recipient strain were mapped to the genome of both CBS 144490 (from which the donor was derived) and for cross-species transfers the corresponding recipient *P. paravariotii* FRR 5287 [32] or *A. fumigatus* A1160 [60] using Bowtie 2 [55]. Coverage over 1 kb non-overlapping windows across the genome was determined using bamCoverage [61]. SNPs were called using FreeBayes [62]. The number of SNPs in each 1 kb window was determined using bedtools intersect [63]. These steps were implemented using a workflow developed on the Australian Galaxy server [53].

The repetitive nature of *Pegasus* and multiple copies of *Hephaestus* limited the use of assembly-based approaches in analyzing the recipient strains. However, the exceptions were the *A. fumigatus* recipient strains #53 and #58 which contained a single copy of *Hephaestus* and no *Pegasus*. The genomes were assembled using Velvet version 1.2.10 [64]. A dot plot to compare the assembly contigs containing *Hephaestus* with the wild type chromosome was generated using D-GENIES [65].

### Phenotyping of metal ion resistances

Resistances to four of the metal ion resistances conferred by *Hephaestus* were assessed on PDA supplemented with CuSO_4_, CdSO_4_ or ZnSO_4_. *Aspergillus* or *Paecilomyces* conidia were suspended in water (10^6^ spores per ml) and 2 μl of this spore suspension was pipetted onto the center of a PDA petri dish (with or without metals added). Strains were incubated for three days (or four days for the plates containing copper sulfate) at 30°C.

### Survey of Paecilomyces Starships

We downloaded 78 publicly available genomes from the *Paecilomyces* genus and combined them with a resequenced long-read assembly of CBS144490 (Table S3.1). Genomes were filtered for contigs smaller than 1kb, soft-masked using Earl Grey ([66]) then annotated using Braker3 ([67]) (--fungus --gff3) using compleasm (--busco_lineage=eurotiales_odb10) and the OrthoDB-11 fungal protein database (--prot_seq). Functional annotation was added using iprscan ([68]), emapper ([69]) and funannotate using the ‘ascomycota’ busco database. All genomes and their resulting proteomes were analysed with BUSCO (eurotiales_odb10). Orthofinder was run on the combined resulting proteomes. For the Paecilomyces genus tree, we used BUSCO (-l eurotiales) [70] was run on all genomes, BUSCOs conserved across all genomes were extracted, aligned using mafft (--auto) [71] and trimmed (-automated1). A species tree was built using iqtree (--alrt 1000 --ufboot 1000 -T AUTO --seed 111111) [72] and the resulting tree was rooted at the midpoint (Figure S6).

We systematically annotated *Starships* in the 79 *Paecilomyces* genomes using starfish v1.1 [35]. First *starfish annotate* was run (-p ∼/starfish/db/YRsuperfams.p1-512.hmm -P ∼/starfish/db/YRsuperfamRefs.faa) to de-novo annotate tyrosine recombinases (tyrR). We combined this annotation with the complete proteome using *starfish consolidate* and then tyrRs were clustered using *starfish sketch*. *Starship* elements were detected using a *starfish insert* sequentially two times, the second time we reduced the conservative default flanking parameters (--pid 80 --hsp 750) to help detection in genomes with larger distances to others. Information on the predicted elements and their associated Captains was summarised using *starfish summarize*. Captains were assigned to families using *hmmsearch* and to different homologous gene groups based on their amino acid sequences using *mmseqs easy-cluster*. *Starships* were then assigned to haplotypes using *starfish sim* and *group*. Each candidate *Starship* was visualised using both *starfish pair-viz* and *element-viz*, then manually inspected. 60/75 candidates with good indicators (large flanking regions for the inserted sequence, haplotype found in multiple unique locations) were kept (Figure S6; Table S3.3). Reference *Starships* of each homologous captain group (navis) and haplotype were designated and manual curation of flanking sequence and insertion sites was conducted to refine the boundaries of each element.

To evaluate the distribution of these *Starships* outside of the *Paecilomyces* genus, the reference sequence for each *Starship* haplotype was BLAST searched against all publicly available NCBI genomes in the Pezizomycotina (10,010 complete genome assemblies downloaded on December 20, 2024). After initially filtering out alignments less than 10 kb and 80% identity, alignments were removed if the total length of the alignment per haplotype was less than 50 kb or covered <50% of the total length of the *Starship* haplotype query. Each alignment was then realigned, with an additional 50 kb added to the distal edges of the most upstream and downstream BLAST alignments, with nucmer [73], the delta file was converted to paf format using paftools *delta2paf* [74] and visualised using gggenomes [75].

### BUSCO phylogeny

We used BUSCO (-l eurotiales) to annotate conserved orthologs in all genomes identified as containing a horizontally transferred *Starship*, plus two additional genomes from the Onygenales (*Blastomyces silverae* (GCA_001014755.1) and *Paracoccidioides brasiliensis* (GCA_000150735.2)) [70]. BUSCOs present in all genomes were extracted, aligned using mafft (--auto) [71] and trimmed (-automated1). A species tree was built using iqtree (--alrt 1000 --ufboot 1000 -T AUTO --seed 111111) using the BUSCO protein sequence alignment [72] and rooted using the two additional Onygenales genomes. We used TimeTree [40] to estimate the timing separation between the *Onygenales*/*Eurotiales* and *Aspergillaceae*/*Thermoascaceae*.

## Data availability

Sequencing data for *P. variotii* CBS 144490 including raw reads and an assembly have been deposited under NCBI BioProject PRJNA1211195. A genome annotation file for CBS 144490 is available on the Melbourne Figshare repository (doi 10.26188/28502201)

## Funding

A.S.U. was supported by the Australian Research Council Discovery Early Career Research Award (DE250100255), a Wenner-Gren Foundation postdoctoral scholarship, and funding from The Royal Physiographic Society of Lund and the Lars Hiertas Minne Foundation. E.G.-T and S.O. are supported by the Office of the Vice Chancellor for Research and Graduate Education at the University of Wisconsin-Madison with funding from the Wisconsin Alumni Research Foundation and the Department of Plant Pathology at the University of Wisconsin-Madison. A.A.V. was provided with support by funding from the Swedish Research Council VR (grant number 2021-04290), and ERC-2023-COG (Starship, 101126121) to A.A.V. Funded by the European Union. Views and opinions expressed are however those of the author(s) only and do not necessarily reflect those of the European Union or the European Research Council Executive Agency. Neither the European Union nor the granting authority can be held responsible for them.

## Acknowledgements

We thank Alexander Idnurm for critical review of the manuscript. We thank Ben Auxier for the insightful discussions. Figure 1A was created in BioRender. https://BioRender.com/b64u749.

## Supplementary results

### *Hephaestus* is integrated into the genome during co-culture via HGT

We assessed the position of *Hephaestus* in each recipient by looking for reads mapping to the flanks of *Hephaestus* and mapping the overhanging portions of these reads back to the acceptor genome (Figure 2C). The obtained sites showed the expected TTAC(N7)A target site previously determined for *Hephaestus* (Figure 2D, Figure S3) [21]. The native copy of *Hephaestus*, if present, was maintained at the expected locus whereas the received copy was integrated elsewhere for the vast majority of pairings. The one exception was *P. paravariotii* FRR 5287 recipient #43 in which the native *Hephaestus* has transposed to a new genomic position. In some cases more integration sites were observed than expected indicating a mixed culture of two recipient strains each containing one new copy obtained during the transfer experiment. For example, in recipient #9 read depth indicated a single additional copy of *Hephaestus* (Figure S1), yet two additional integration sites were observed (Figure S3).

### Characterisation of the *Pegasus Starship*

Given that *Pegasus* had transferred in our experiment we further investigated the biological features of this transposon. In the previously available assembly of *P. variotii* CBS 144490, *Pegasus* was fragmented across multiple assembly contigs, likely due to internal duplications [21]. In the only other genome known to contain *Pegasus*, that of *Paecilomyces lecythidis* strain MCCF 102, *Pegasus* is highly fragmented and has been inactivated through mutation [21]. Thus the full sequence of *Pegasus* was unknown and it was not known if the CBS 144490 copy of *Pegasus* was an active *Starship*. To determine the complete sequence of *Pegasus* we resequenced the CBS 144490 genome with Oxford Nanopore technology and used these long-reads to assemble the genome into ten nuclear contigs and one mitochondrial contig. Within this assembly *Pegasus* was completely assembled on contig 2. *Pegasus* is 403.6 kb and it contains 123 predicted genes (lifted over from our previous annotated assembly [34]) including characteristic *Starship* genes, namely a captain recombinase, and two DUF3723 domain proteins. The function of DUF3723 domain proteins is not known but they are commonly found in Starships [22]. In addition it contains a further captain recombinase and DUF3723 belonging to a potentially nested *Starship* (Figure S4A). Similar to previous work on *Hephaestus* we generated a mini-element including the terminal sequences, the captain and a selectable marker, and embedded this mini-element within a hygromycin-resistance cassette (Figure S4B). This allowed for us to screen for transposition of the mini-element. By sequencing a pool of strains in which *Pegasus* had transposed we were able to identify 47 insertion sites of mini-*Pegasus* which were broadly distributed across the genome (Figure S4C). These insertion sites revealed a 10 bp palindromic target site TACCTAGGTA (Figure S4D).

### Survey of *Starships* in *Paecilomyces*

We combined both automated and manual annotations of *Starships* across the *Paecilomyces* genus using a set of 79 genomes, spanning 10 phylogenetically validated species (Figure S6; Table S3.1; Table S3.2). We detected 33 distinct *Starship* haplotypes from 6 Captain tyrR families (*Enterprise, Galactica, Hephaestus, Phoenix, Prometheus* and *Tardis*) ranging from 16-403 kb in size (Figure S6; Table S3.3).

We then systematically searched for recently transferred copies of these elements using publicly available NCBI genome assemblies for the Pezizomycotina (n=10,010), the largest subphylum of filamentous fungi predicted to be the primary carriers of *Starships* (Table S3.1) [35]. Initially, to support the assertion that highly similar regions observed among different species are likely a result of HGT, we show that the ability to align genomic sequences is rapidly lost within the *Paecilomyces* genus when comparing increasingly more distantly related species, using the long-read assembly of *P. formosus* as a reference (Figure S16). Pairwise whole-genome alignments to the long-read *P. formosus* assembly covered 12-96% of the genome with average alignment lengths of 1-30 kb across all genome-genome comparisons within *Paecilomyces*.

This reduced further to <1% coverage and <1 kb alignments on average when aligning genomes from other genera to the *P. formosus* assembly, emphasizing a drastic decrease in the recovery of highly similar and large sequences by chance (Figure S16). The species IDs of all putative donors and recipients were also confirmed by building a BUSCO-based maximum likelihood phylogeny (Figure 3; Table S3.2).

Pairwise alignments between each *Starship* copy and the genomes in which they were identified revealed an average identity of 88-99% and an average total alignment length of 52-286 kb (Figure S16). Notably, the average pairwise nucleotide similarity of all *Starships/*genomic regions identified as originating from *Keraunos* was 93% across 13 species spanning four genera and two taxonomic orders, the *Eurotiales* and *Onygenales*.

Additionally, we now have evidence that two different *Starships* have been transferred between different strains of *Paecilomyces variotii* and *Spiromastigoides* sp. These two *Starships* contain captains from different families and do not align to one another. This indicates that the movement of *Starships* over such long distances is not restricted to particular Captain tyrR families and is a more general feature of *Starship* elements. The alignment of *Keraunos* elements from different species further indicated that a tandem insertion of a different *Starship* at the 5’ end of a subset of *Keraunos* elements has created a hybrid *Starship* that appears to horizontally transfer as a single contiguous element. The captain of this additional *Starship* is closely related to captain tyrRs from the previously described *Mithridate Starship* (Supp figure of captain tree?) [29]. The transfer of *Hera* appears to be a potential case of cargo-swapping, as alignment to the captain tyrR in the reference element was missing in all cases of identified similarity with other genomes (Figure S13). 2 out of the 9 ships with evidence of HGT contain a MYB/SANT-like transcription factor at the edge of the element opposite the captain (Figure S11; Figure S12) providing more evidence for the conserved structural position of this gene family in particular clades of *Starship* elements [28].

## Supplementary discussion

### Why HGT is the most likely explanation to explain the outcomes of the experiments

One alternative to HGT is parasexual recombination. While not specifically reported in *Paecilomyces,* parasex has been long known in a number of better-studied *Eurotiales* species and it is very possible that *Paecilomyces* may undergo this process [76,77]. This may be a particularly compelling explanation for the *Starship* transfer to an isogenic *P. variotii* acceptor. However, the expected result of parasex would be the introduction of *Hephaestus* carrying hygromycin resistance at the parent locus, ie, replacing the wild type *Hephaestus* copy of the acceptor strain through recombination. But what we observed in the Illumina sequencing was an increase in sequencing coverage over *Hephaestus* and the clean integration of one or more *Hephaesti* at new TTACN_7_A target sites within the genome.

A related explanation for the observed *Starship* transfer in the isogenic pairing would be the transposition of *Hephaestus* (and additionally where relevant, *Pegasus*) within the donor genome followed by parasexual recombination to result in strains having two copies. While theoretically possible, such an explanation would beg the question as to why we did not observe the other expected outcomes of this parasexual event, for example donor isolates that have inherited the *leuA* gene of the acceptor (which would complement the donor’s leucine-deficient phenotype). Furthermore, parasexual recombination is usually limited to within species and there is no precedent to our knowledge of parasex between genera, which would be required to explain the transfer into *Aspergillus*. Moreover, in the observed interspecies transfers, the genetic divergence between the donor and acceptor genomes allows us to determine that the only donor-derived DNA in the recipient strains is the *Starship* DNA. This would be a highly unlikely outcome from parasex alone.

Another alternative explanation might be that the apparent recipient strains were in fact a heterokaryon or diploid between the donor and the recipient strain. While interspecies, and even intergenus heterokaryons have been reported, these are typically created under highly artificial conditions such as protoplast fusion and not through simple co-culture where vegetative incompatibility is expected to block fusion [78]. Additionally, in a heterokaryon we would expect to see additional DNA other than the *Starship* derived from the donor present in the recipient and would expect the apparent copy number of the transferred *Starship* to be less than one because not all nuclei in the heterokaryon would contain a copy of the *Starship*.

### Horizontal Chromosome Transfer: a special case of parasex?

Previous work has observed the horizontal transfer of entire chromosomes in fungal co-cultures. However, these experimental studies are limited to transfers between closely related strains within a species or species complex [79–84]. It has been suggested that such transfers might be the result of parasex followed by selective chromosome inheritance [84]. However, intriguing alternative hypotheses independent of karyogamy have been proposed such as preferential transfer between nuclei [85]. A reliance on parasex would explain why such transfers appear to be restricted to short phylogenetic distances, typically within a single species. The only interspecific HCT to be discovered, between *Metarhizium guizhouense* and *Metarhizium robertsii,* involves two species separated by only 2.3 MY - 15.1 MY, far less than found here for *Starships* [86][83]. However, in other genera such as *Cryptococcus,* hybridisation remains possible across divergence times as large as what we observed [87]. We believe that further work is needed to establish HCT as a means of genetic exchange in fungi separate to parasex.

## Supplemental Tables and Figures

**Table S1:** Outcomes of co-culture experiments between the *P. variotii* donor and 3 acceptor strains.

| Plate number | Acceptor strain | Recipient produced? | Verification |
| --- | --- | --- | --- |
| 1 | *P. variotii* CBS 144490 | yes | Illumina |
| 2 | *P. variotii* CBS 144490 | yes | Illumina |
| 3 | *P. variotii* CBS 144490 | yes | PCR |
| 4 | *P. variotii* CBS 144490 | yes | PCR |
| 37 | *P. variotii* CBS 144490 | yes | Illumina |
| 38 | *P. variotii* CBS 144490 | no | N/A |
| 39 | *P. variotii* CBS 144490 | no | N/A |
| 40 | *P. variotii* CBS 144490 | yes | Illumina |
| 41 | *P. variotii* CBS 144490 | no | N/A |
| 42 | *P. variotii* CBS 144490 | no | N/A |
| 43 | *P. variotii* CBS 144490 | yes | Illumina |
| 44 | *P. variotii* CBS 144490 | yes | Illumina |
| 45 | *P. variotii* CBS 144490 | yes | Illumina |
| 46 | *P. variotii* CBS 144490 | no | N/A |
| 47 | *P. variotii* CBS 144490 | yes | Illumina |
| 48 | *P. variotii* CBS 144490 | no | N/A |
| 5 | *P. paravariotii* FRR 5287 | no | N/A |
| 6 | *P. paravariotii* FRR 5287 | no | N/A |
| 7 | *P. paravariotii* FRR 5287 | yes | Illumina |
| 8 | *P. paravariotii* FRR 5287 | yes | Illumina |
| 9 | *P. paravariotii* FRR 5287 | yes | Illumina |
| 10 | *P. paravariotii* FRR 5287 | yes | Illumina |
| 11 | *P. paravariotii* FRR 5287 | yes | Illumina |
| 12 | *P. paravariotii* FRR 5287 | yes | Illumina |
| 13 | *P. paravariotii* FRR 5287 | yes | Illumina |
| 14 | *P. paravariotii* FRR 5287 | yes | Illumina |
| 15 | *P. paravariotii* FRR 5287 | yes | Illumina |
| 16 | *P. paravariotii* FRR 5287 | yes | Illumina |
| 17 | *P. paravariotii* FRR 5287 | yes | Illumina |
| 18 | *P. paravariotii* FRR 5287 | yes | Illumina |
| 19 | *P. paravariotii* FRR 5287 | no | N/A |
| 20 | *P. paravariotii* FRR 5287 | yes | Illumina |
| 21 | *P. paravariotii* FRR 5287 | yes | Illumina |
| 22 | *P. paravariotii* FRR 5287 | yes | Illumina |
| 59 | *P. paravariotii* FRR 5287 | yes | PCR |
| 60 | *P. paravariotii* FRR 5287 | yes | PCR |
| 61 | *P. paravariotii* FRR 5287 | yes | PCR |
| 62 | *P. paravariotii* FRR 5287 | yes | PCR |
| 63 | *P. paravariotii* FRR 5287 | yes | - |
| 64 | *P. paravariotii* FRR 5287 | yes | PCR |
| 65 | *P. paravariotii* FRR 5287 | yes | PCR |
| 66 | *P. paravariotii* FRR 5287 | yes | PCR |
| 67 | *P. paravariotii* FRR 5287 | yes | PCR |
| 68 | *P. paravariotii* FRR 5287 | yes | PCR |
| 69 | *P. paravariotii* FRR 5287 | yes | PCR |
| 70 | *P. paravariotii* FRR 5287 | yes | PCR |
| 71 | *P. paravariotii* FRR 5287 | yes | PCR |
| 72 | *P. paravariotii* FRR 5287 | yes | PCR |
| 73 | *P. paravariotii* FRR 5287 | yes | PCR |
| 74 | *P. paravariotii* FRR 5287 | yes | PCR |
| 75 | *P. paravariotii* FRR 5287 | yes | PCR |
| 76 | *P. paravariotii* FRR 5287 | yes | PCR |
| 77 | *P. paravariotii* FRR 5287 | yes | PCR |
| 78 | *P. paravariotii* FRR 5287 | yes | PCR |
| 79 | *P. paravariotii* FRR 5287 | yes | PCR |
| 80 | *P. paravariotii* FRR 5287 | yes | PCR |
| 81 | *P. paravariotii* FRR 5287 | yes | PCR |
| 82 | *P. paravariotii* FRR 5287 | yes | PCR |
| 83 | *P. paravariotii* FRR 5287 | yes | PCR |
| 84 | *P. paravariotii* FRR 5287 | yes | PCR |
| 85 | *P. paravariotii* FRR 5287 | yes | PCR |
| 86 | *P. paravariotii* FRR 5287 | yes | PCR |
| 87 | *P. paravariotii* FRR 5287 | yes | PCR |
| 88 | *P. paravariotii* FRR 5287 | yes | PCR |
| 89 | *P. paravariotii* FRR 5287 | yes | PCR |
| 90 | *P. paravariotii* FRR 5287 | yes | PCR |
| 91 | *P. paravariotii* FRR 5287 | yes | PCR |
| 92 | *P. paravariotii* FRR 5287 | yes | PCR |
| 93 | *P. paravariotii* FRR 5287 | yes | PCR |
| 94 | *P. paravariotii* FRR 5287 | yes | PCR |
| 95 | *P. paravariotii* FRR 5287 | yes | PCR |
| 96 | *P. paravariotii* FRR 5287 | yes | PCR |
| 97 | *P. paravariotii* FRR 5287 | yes | PCR |
| 98 | *P. paravariotii* FRR 5287 | yes | PCR |
| 99 | *P. paravariotii* FRR 5287 | yes | PCR |
| 100 | *P. paravariotii* FRR 5287 | yes | PCR |
| 101 | *P. paravariotii* FRR 5287 | yes | PCR |
| 102 | *P. paravariotii* FRR 5287 | yes | PCR |
| 103 | *P. paravariotii* FRR 5287 | yes | - |
| 104 | *P. paravariotii* FRR 5287 | yes | PCR |
| 105 | *P. paravariotii* FRR 5287 | yes | PCR |
| 106 | *P. paravariotii* FRR 5287 | yes | PCR |
| 107 | *P. paravariotii* FRR 5287 | yes | PCR |
| 108 | *P. paravariotii* FRR 5287 | yes | PCR |
| 109 | *P. paravariotii* FRR 5287 | yes | PCR |
| 110 | *P. paravariotii* FRR 5287 | yes | PCR |
| 111 | *P. paravariotii* FRR 5287 | yes | PCR |
| 112 | *P. paravariotii* FRR 5287 | yes | PCR |
| 113 | *P. paravariotii* FRR 5287 | yes | PCR |
| 114 | *P. paravariotii* FRR 5287 | yes | PCR |
| 115 | *P. paravariotii* FRR 5287 | yes | PCR |
| 116 | *P. paravariotii* FRR 5287 | yes | PCR |
| 117 | *P. paravariotii* FRR 5287 | yes | PCR |
| 118 | *P. paravariotii* FRR 5287 | yes | PCR |
| 119 | *P. paravariotii* FRR 5287 | yes | PCR |
| 120 | *P. paravariotii* FRR 5287 | yes | PCR |
| 121 | *P. paravariotii* FRR 5287 | yes | PCR |
| 122 | *P. paravariotii* FRR 5287 | yes | PCR |
| 123 | *P. paravariotii* FRR 5287 | yes | PCR |
| 124 | *P. paravariotii* FRR 5287 | yes | PCR |
| 125 | *P. paravariotii* FRR 5287 | yes | PCR |
| 126 | *P. paravariotii* FRR 5287 | yes | PCR |
| 127 | *P. paravariotii* FRR 5287 | yes | PCR |
| 128 | *P. paravariotii* FRR 5287 | yes | PCR |
| 129 | *P. paravariotii* FRR 5287 | yes | PCR |
| 130 | *P. paravariotii* FRR 5287 | yes | PCR |
| 131 | *P. paravariotii* FRR 5287 | yes | PCR |
| 132 | *P. paravariotii* FRR 5287 | yes | PCR |
| 133 | *P. paravariotii* FRR 5287 | yes | PCR |
| 134 | *P. paravariotii* FRR 5287 | yes | PCR |
| 135 | *P. paravariotii* FRR 5287 | yes | PCR |
| 136 | *P. paravariotii* FRR 5287 | yes | PCR |
| 137 | *P. paravariotii* FRR 5287 | yes | PCR |
| 138 | *P. paravariotii* FRR 5287 | yes | PCR |
| 23 | *A. fumigatus* A1160 | no | N/A |
| 24 | *A. fumigatus* A1160 | no | N/A |
| 25 | *A. fumigatus* A1160 | no | N/A |
| 26 | *A. fumigatus* A1160 | no | N/A |
| 27 | *A. fumigatus* A1160 | yes | Illumina |
| 28 | *A. fumigatus* A1160 | no | N/A |
| 29 | *A. fumigatus* A1160 | no | N/A |
| 30 | *A. fumigatus* A1160 | no | N/A |
| 31 | *A. fumigatus* A1160 | no | N/A |
| 32 | *A. fumigatus* A1160 | no | N/A |
| 33 | *A. fumigatus* A1160 | no | N/A |
| 34 | *A. fumigatus* A1160 | no | N/A |
| 35 | *A. fumigatus* A1160 | no | N/A |
| 36 | *A. fumigatus* A1160 | no | N/A |
| 49 | *A. fumigatus* A1160 | no | N/A |
| 50 | *A. fumigatus* A1160 | no | N/A |
| 51 | *A. fumigatus* A1160 | no | N/A |
| 52 | *A. fumigatus* A1160 | no | N/A |
| 53 | *A. fumigatus* A1160 | yes | Illumina |
| 54 | *A. fumigatus* A1160 | no | N/A |
| 56 | *A. fumigatus* A1160 | no | N/A |
| 57 | *A. fumigatus* A1160 | no | N/A |
| 58 | *A. fumigatus* A1160 | yes | Illumina |

**Table S2:** Genotypes of strains sequenced with Illumina.

| Sequenced strain | Acceptor strain | Copy number |  | SRA accession |
| --- | --- | --- | --- | --- |
|  |  | *Pegasus* | *Hephaestus* |  |
| *P. variotii* CBS 144490 | N/A | 1 | 1 |  |
| *P. paravariotii* FRR 5287 | N/A | 0 | 1 |  |
| *A. fumigatus* A1160 | N/A | 0 | 0 |  |
| Donor | N/A | 1 | 1 |  |
| Recipient number 1 | *P. variotii* CBS 144490 | 1 | 2 |  |
| Recipient number 2 | *P. variotii* CBS 144490 | 1 | 2 |  |
| Recipient number 7 | *P. paravariotii* FRR 5287 | 1 | 2 |  |
| Recipient number 8 | *P. paravariotii* FRR 5287 | 1 | 2 |  |
| Recipient number 9 | *P. paravariotii* FRR 5287 | 1 | 2 |  |
| Recipient number 10 | *P. paravariotii* FRR 5287 | 1 | 2 |  |
| Recipient number 11 | *P. paravariotii* FRR 5287 | 2 | 2 |  |
| Recipient number 12 | *P. paravariotii* FRR 5287 | 1 | 2 |  |
| Recipient number 13 | *P. paravariotii* FRR 5287 | 1 | 2 |  |
| Recipient number 14 | *P. paravariotii* FRR 5287 | 1 | 2 |  |
| Recipient number 15 | *P. paravariotii* FRR 5287 | 2 | 2 |  |
| Recipient number 16 | *P. paravariotii* FRR 5287 | 1 | 2 |  |
| Recipient number 17 | *P. paravariotii* FRR 5287 | 2 | 2 |  |
| Recipient number 18 | *P. paravariotii* FRR 5287 | 1 | 2 |  |
| Recipient number 20 | *P. paravariotii* FRR 5287 | 2 | 2 |  |
| Recipient number 21 | *P. paravariotii* FRR 5287 | 1 | 2 |  |
| Recipient number 22 | *P. paravariotii* FRR 5287 | 1 | 2 |  |
| Recipient number 27 | *A. fumigatus* A1160 | 0 | 2 |  |
| Recipient number 37 | *P. variotii* CBS 144490 | 2 | 2 |  |
| Recipient number 40 | *P. variotii* CBS 144490 | 1 | 2 |  |
| Recipient number 43 | *P. variotii* CBS 144490 | 2 | 2 |  |
| Recipient number 44 | *P. variotii* CBS 144490 | 2 | 3 |  |
| Recipient number 45 | *P. variotii* CBS 144490 | 1 | 3 |  |
| Recipient number 47 | *P. variotii* CBS 144490 | 2 | 2 |  |
| Recipient number 53 | *A. fumigatus* A1160 | 0 | 1 |  |
| Recipient number 58 | *A. fumigatus* A1160 | 0 | 1 |  |

**Table S4:**
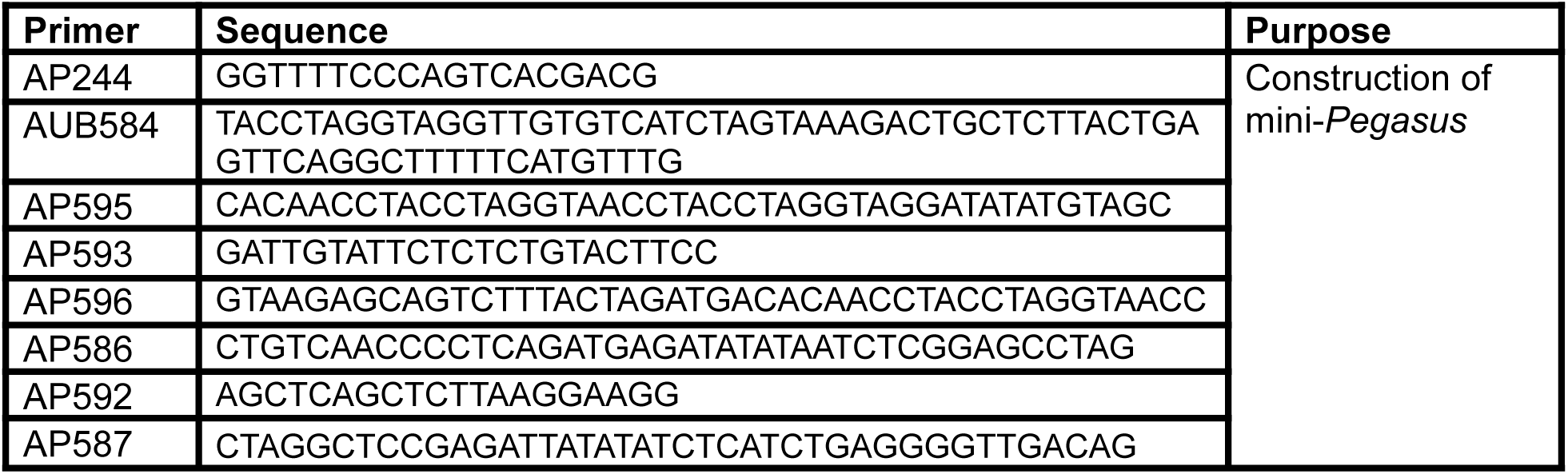
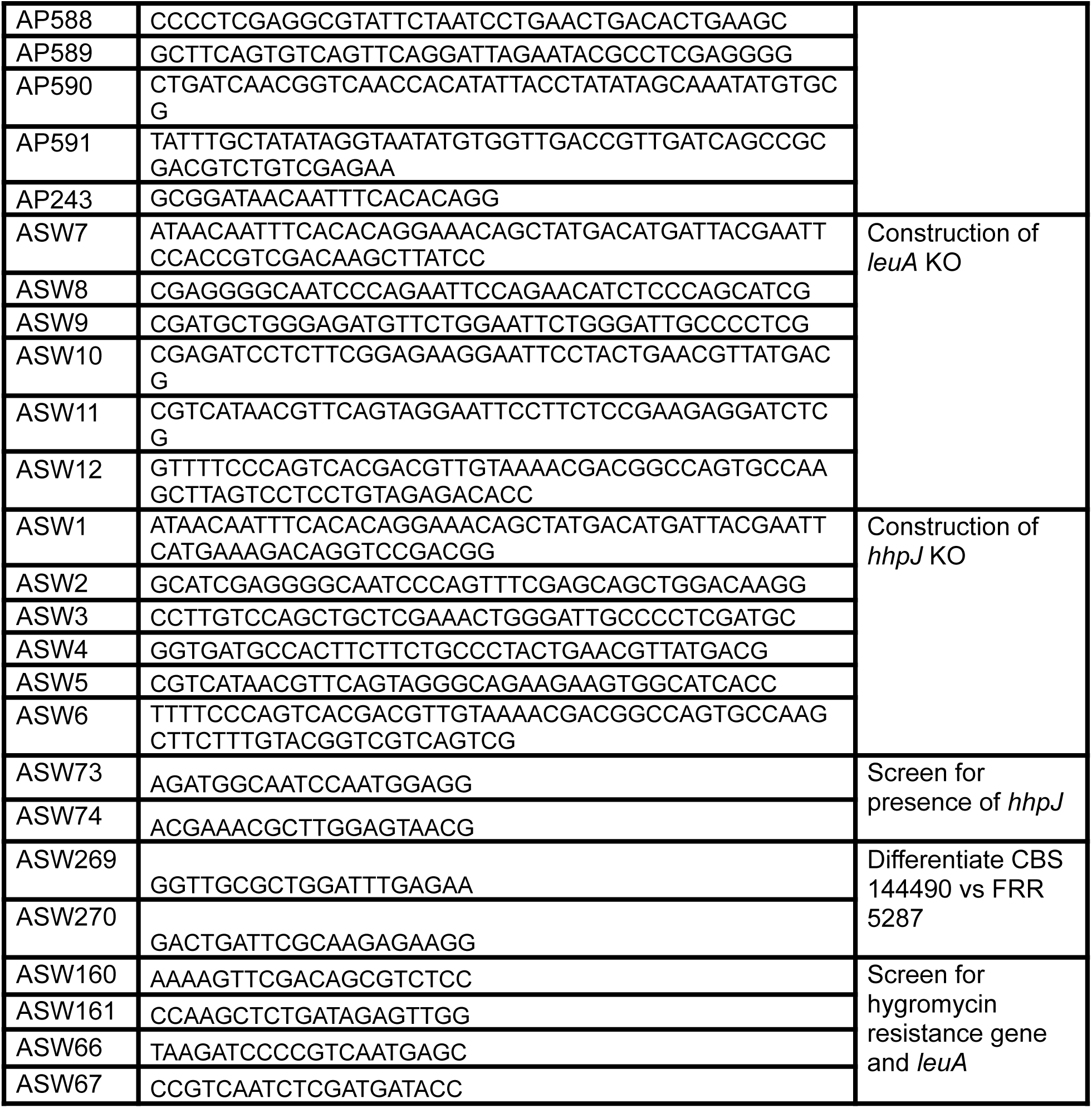
Primers used in this study.

**Figure S1:**
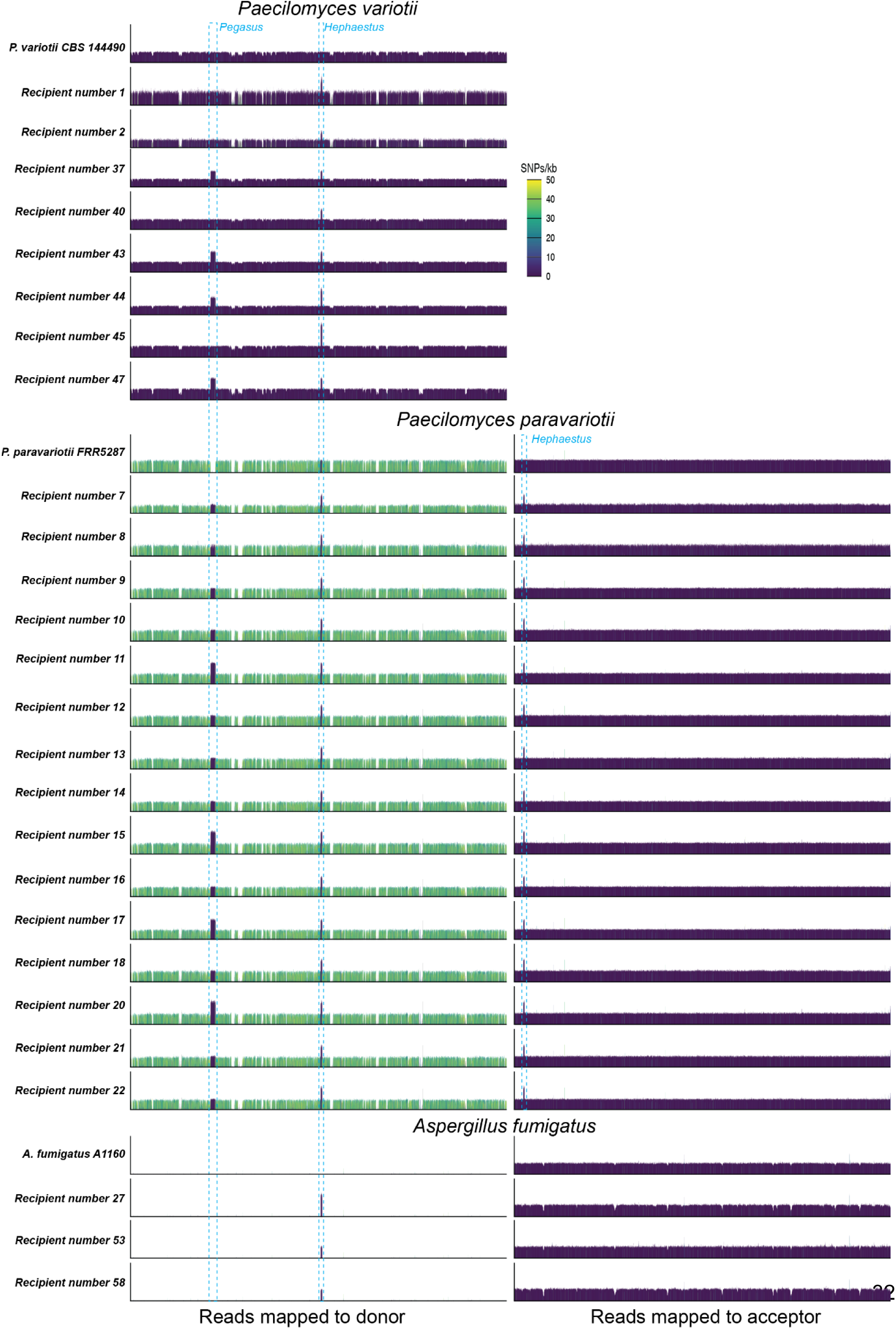
Illumina reads from recipient and wild type strains mapped to donor and acceptor genomes. Assembly contigs were concatenated prior to read mapping.

**Figure S2:**
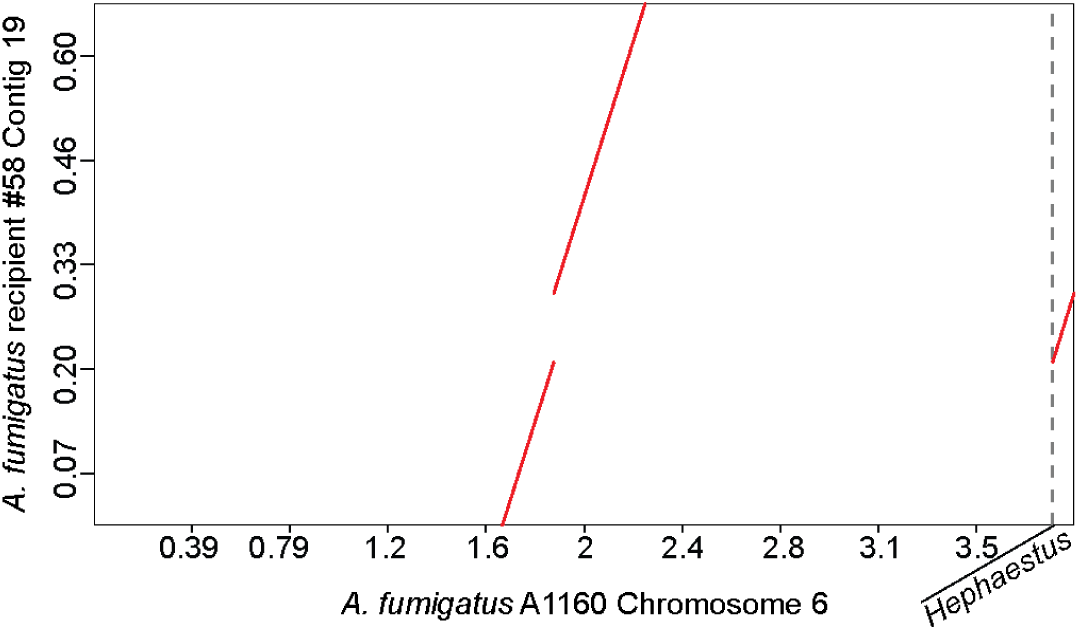
Dot plot comparison between the assembly contig of recipient #58 which contains *Hephaestus* and the corresponding wild type *A. fumigatus* A1160 chromosome.

**Figure S3:**
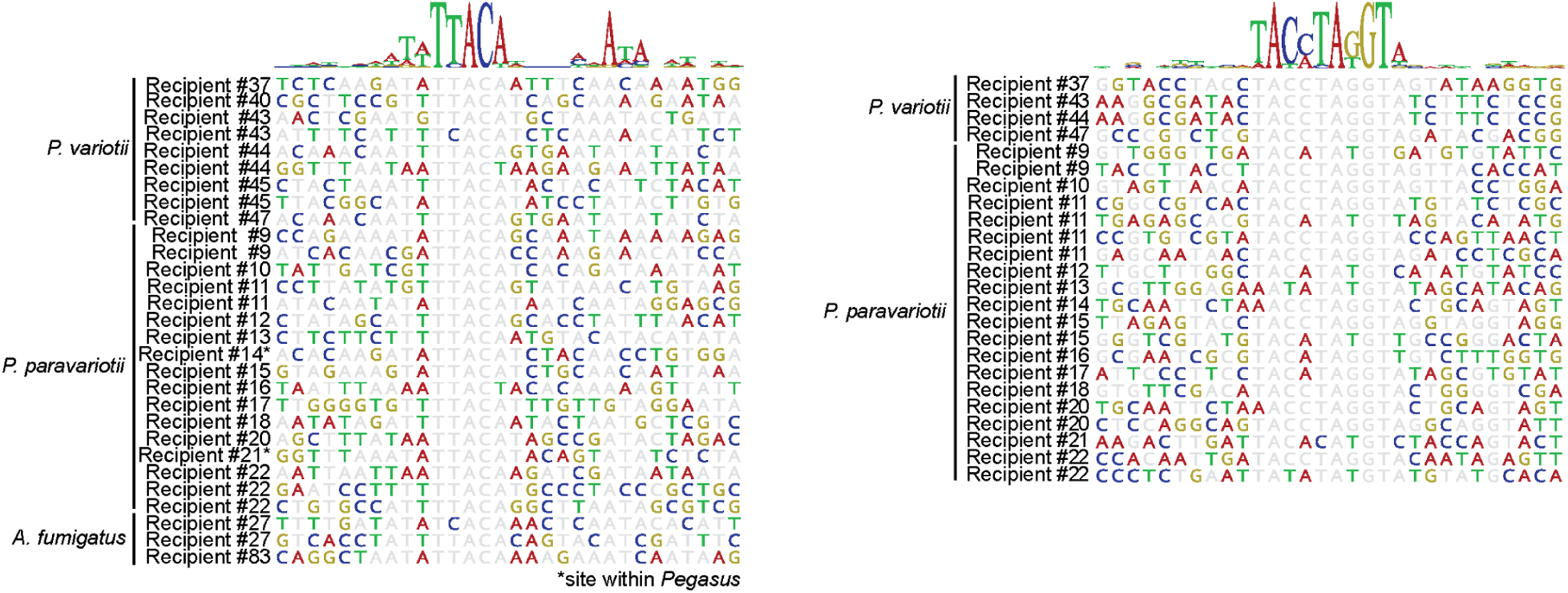
Integration sites of *Hephaestus* and *Pegasus* within recipient genomes.

**Figure S4:**
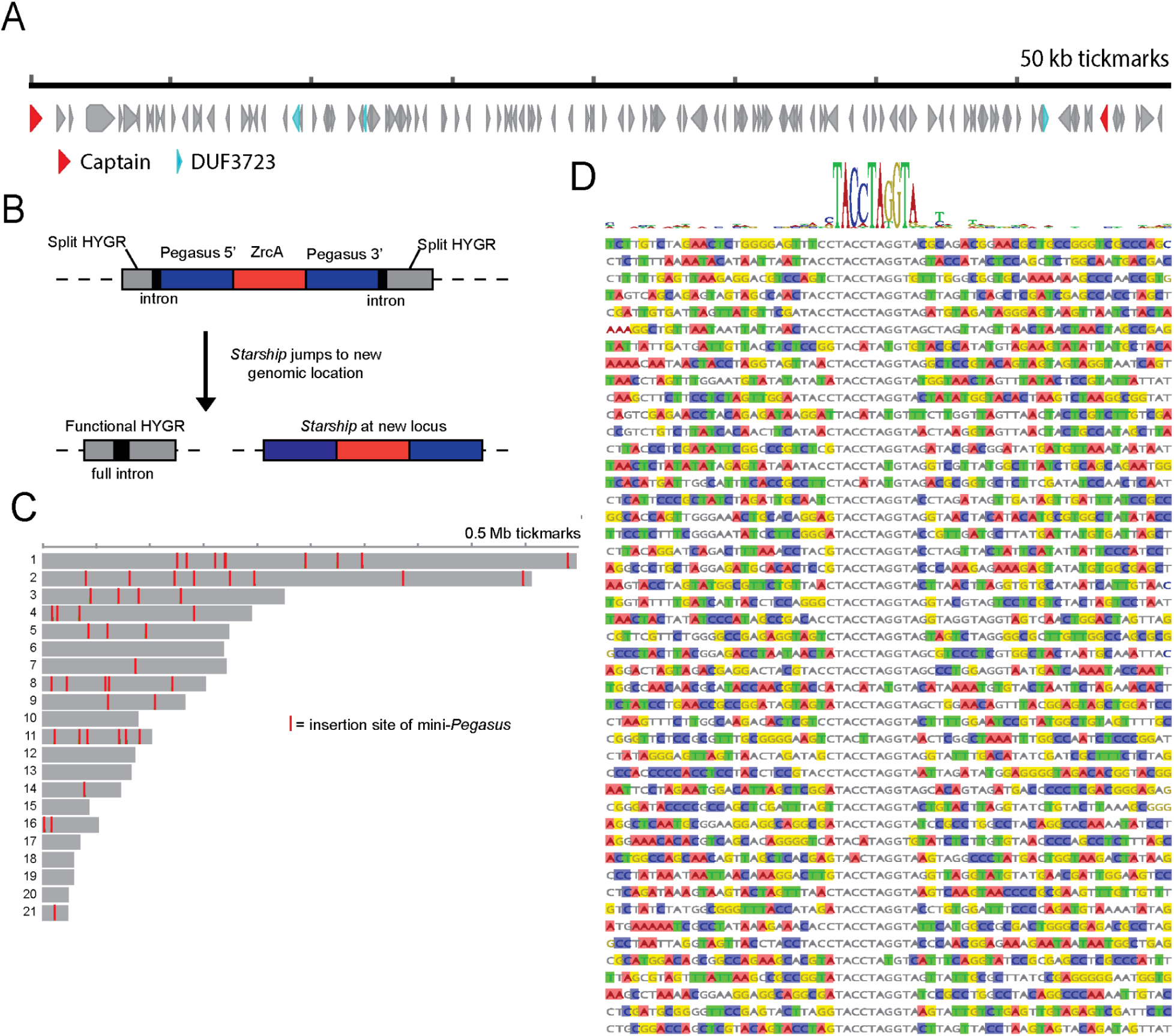
***Pegasus* is an active *Starship* within the *Paecilomyces variotii* genome. A)** gene map of the completely assembled *Pegasus Starship* showing captain and DUF3723 domain genes. **B)** The approach used to demonstrate the transposition of a minimal version of *Pegasus* embedded within a marker for hygromycin resistance. **C)** Insertion sites of minimal *Pegasus* across the genome. Shown are the largest 21 contigs which represent 94% of the complete assembly. **D)** Sequence alignment of mini-*Pegasus* insertion sites with sequence logo.

**Figure S5:**
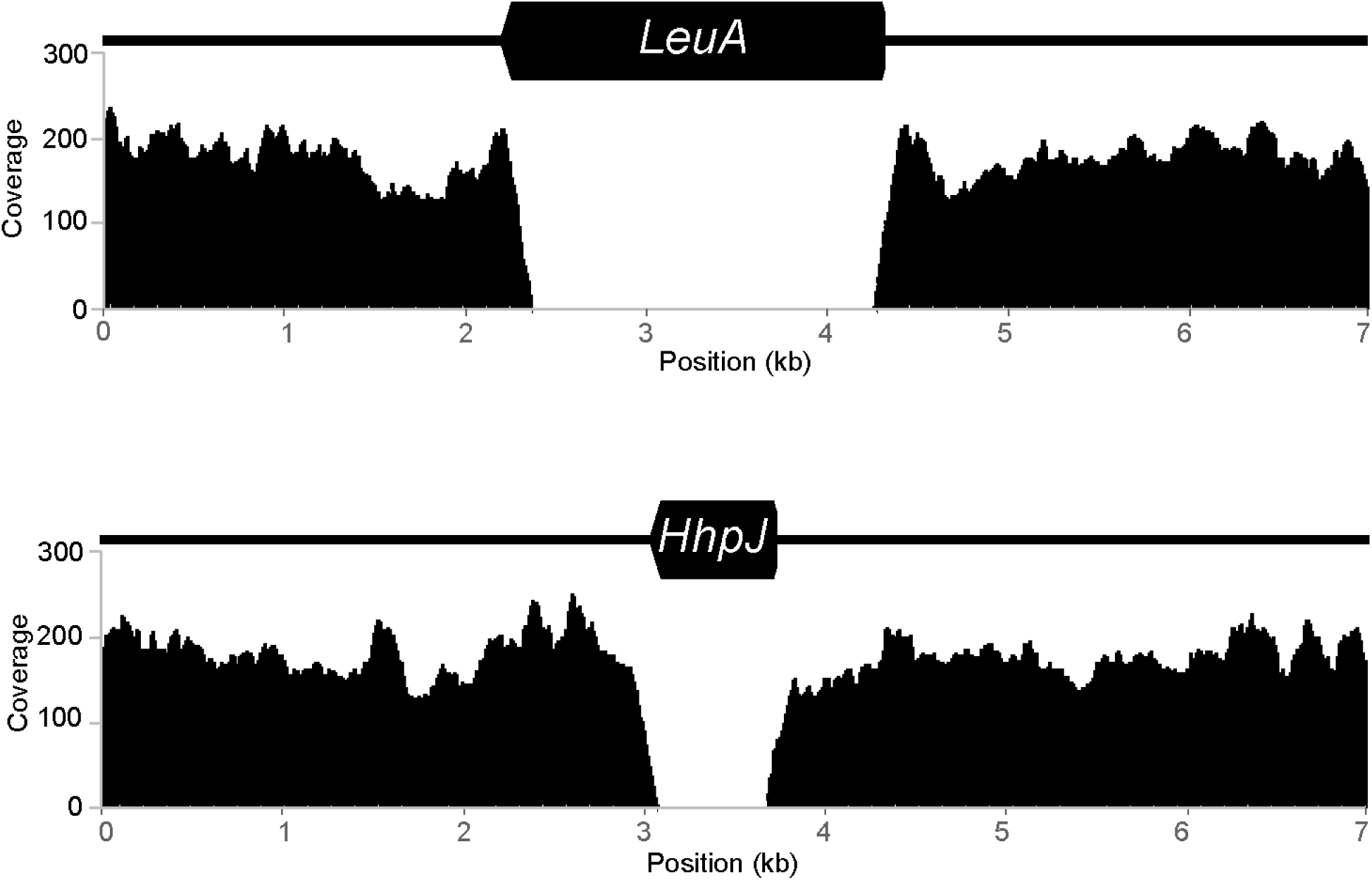
Mapping density of Illumina reads derived from the donor strain to the wild type *leuA* and *hhpJ* gene regions.

**Figure S6:**
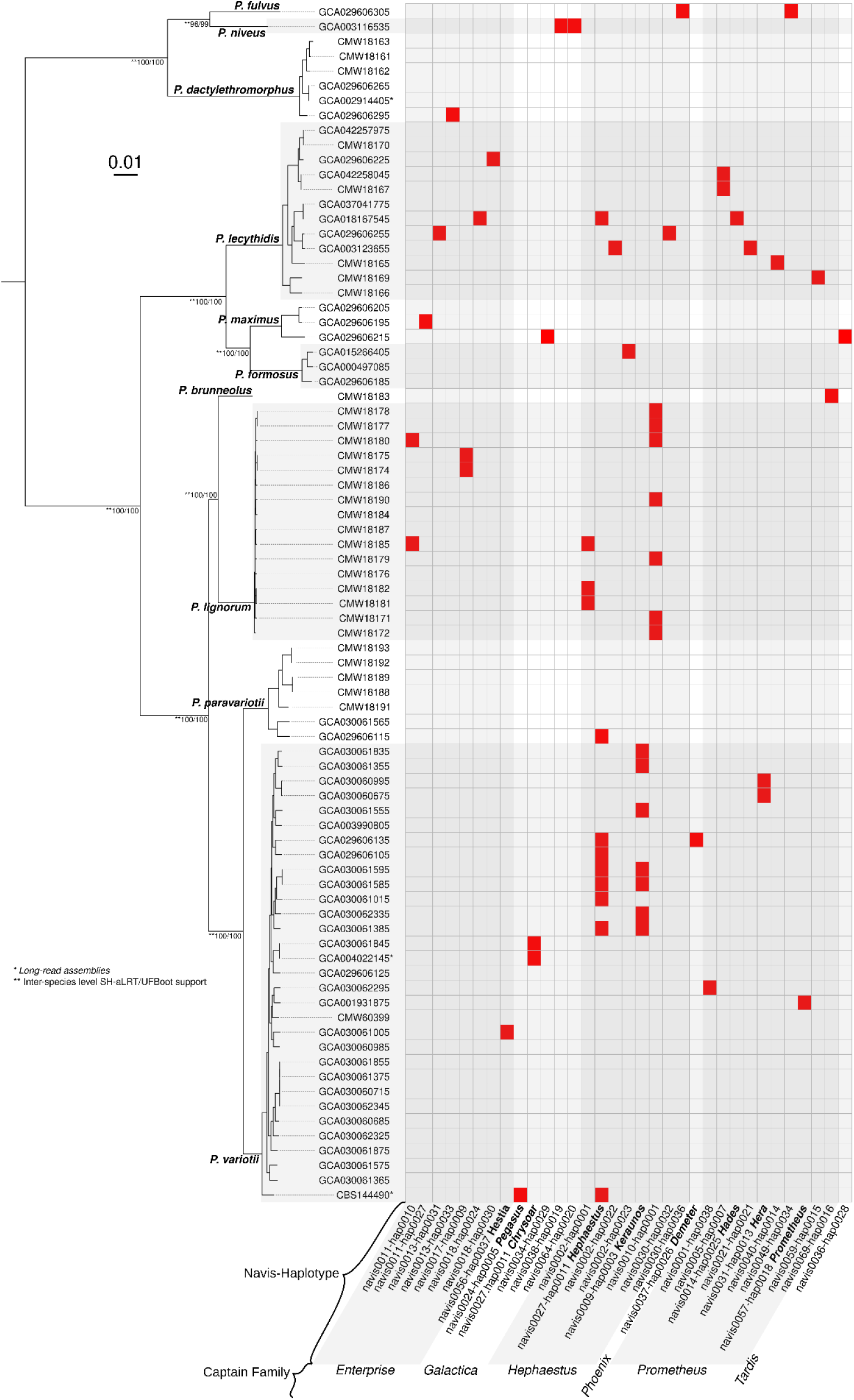
Distribution of 33 high-quality *Starships* annotated with starfish v.1.1 within 79 isolates across the *Paecilomyces* genus. Presence (red) and absence (gray/white) shown in heatmap to the right. The maximum likelihood phylogeny (left) was generated using conserved BUSCO orthologs from the 79 *Paecilomyces* genomes and is midpoint rooted. Ultra-fast bootstrap and aLRT support percentages (out of 1000) are shown below each branch. Tips are labeled by accession.

**Figure S7:**
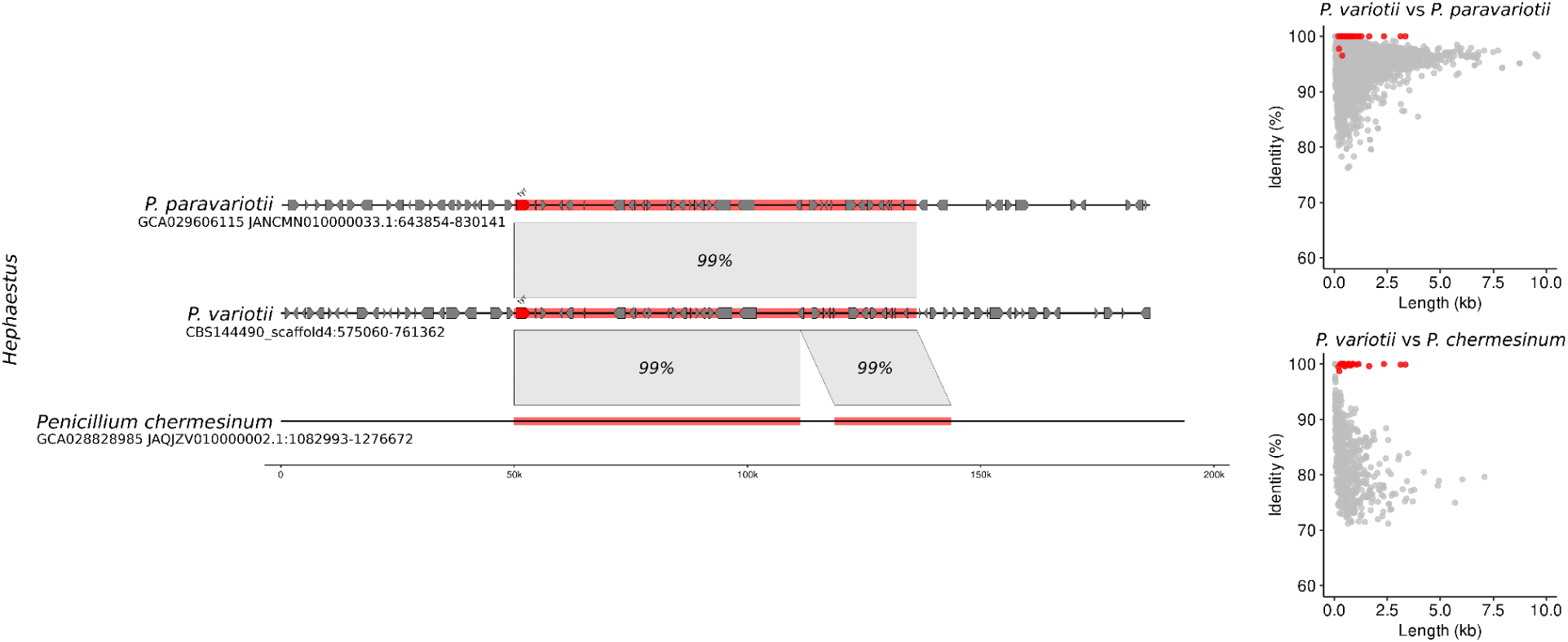
Synteny alignments (left) for the *Hephaestus Starship* shown for species with evidence of its horizontal transfer. Predicted genes are depicted as arrows. *De novo* gene prediction was performed only for *Paecilomyces* isolates. BLASTall plots (right) showing nucleotide BLAST alignment length and identity using all *P. variotii* protein-coding gene sequences as queries and whole genome sequences from the other named isolates as targets. Predicted genes from *Hephaestus* are shown in red.

**Figure S8:**
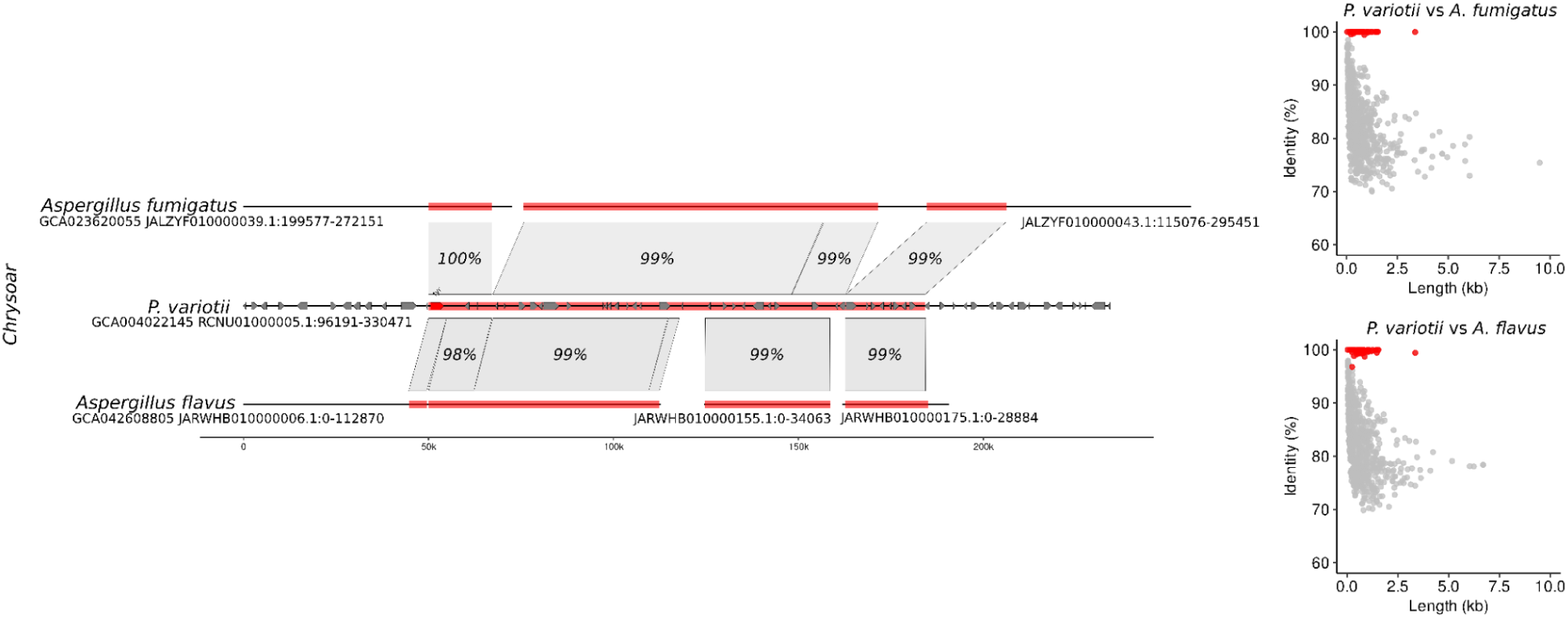
Synteny alignments (left) for the *Chrysaor Starship* shown for species with evidence of its horizontal transfer. Predicted genes are depicted as arrows. *De novo* gene prediction was performed only for *Paecilomyces* isolates. BLASTall plots (right) showing nucleotide BLAST alignment length and identity using all *P. variotii* protein-coding gene sequences as queries and whole genome sequences from the other named isolates as targets. Predicted genes from *Chrysaor* are shown in red.

**Figure S9:**
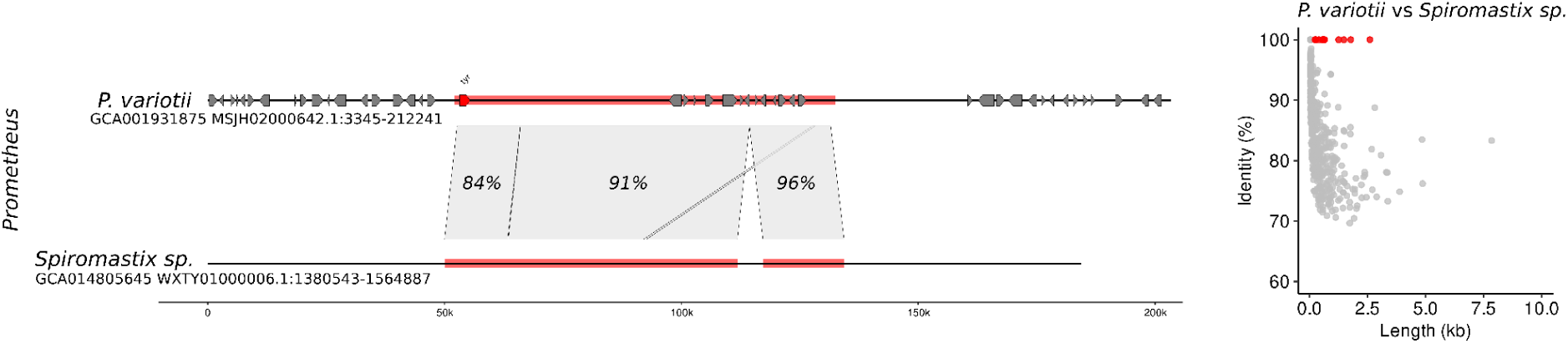
Synteny alignments (left) for the *Prometheus Starship* shown for species with evidence of its horizontal transfer. Predicted genes are depicted as arrows. *De novo* gene prediction was performed only for *Paecilomyces* isolates. BLASTall plots (right) showing nucleotide BLAST alignment length and identity using *P. variotii* protein-coding gene sequences as queries and whole genome sequence from *Spiromastix* as a target. Predicted genes from *Prometheus* are shown in red.

**Figure S10:**
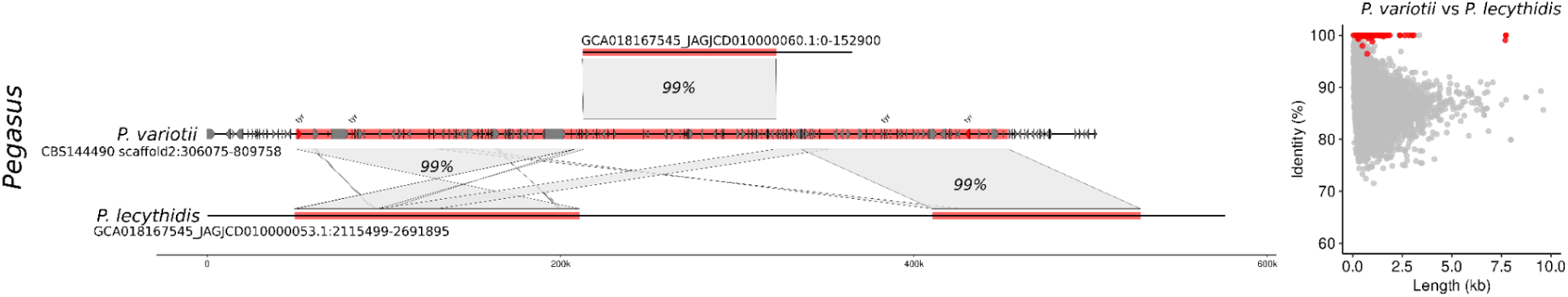
Synteny alignments (left) for the *Pegasus Starship* shown for species with evidence of its horizontal transfer. Predicted genes are depicted as arrows. *De novo* gene prediction was performed only for *Paecilomyces* isolates. BLASTall plots (right) showing nucleotide BLAST alignment length and identity using *P. variotii* protein-coding gene sequences as queries and whole genome sequence from *P. lecythidis* as a target. Predicted genes from *Pegasus* are shown in red.

**Figure S11:**
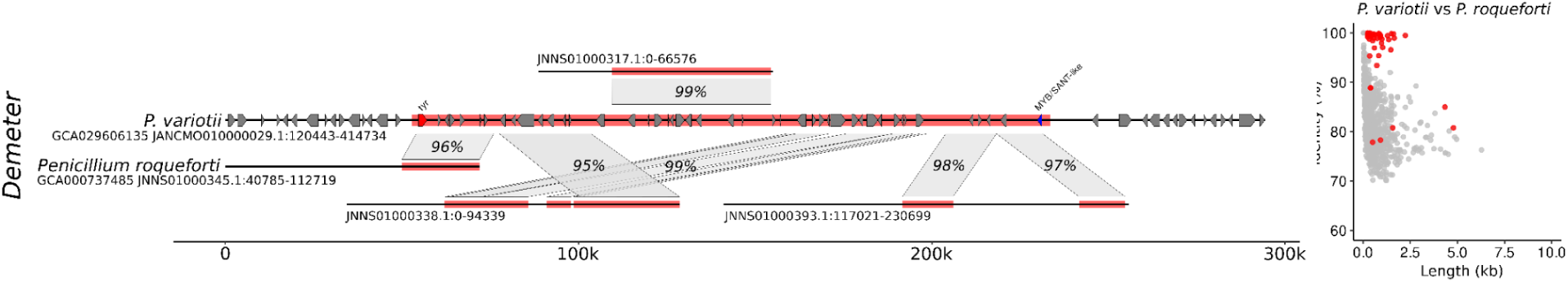
Synteny alignments (left) for the *Demeter Starship* shown for species with evidence of its horizontal transfer. Predicted genes are depicted as arrows. *De novo* gene prediction was performed only for *Paecilomyces* isolates. BLASTall plots (right) showing nucleotide BLAST alignment length and identity using *P. variotii* protein-coding gene sequences as queries and whole genome sequence from *P. roquefortii* as a target. Predicted genes from *Demeter* are shown in red.

**Figure S12:**
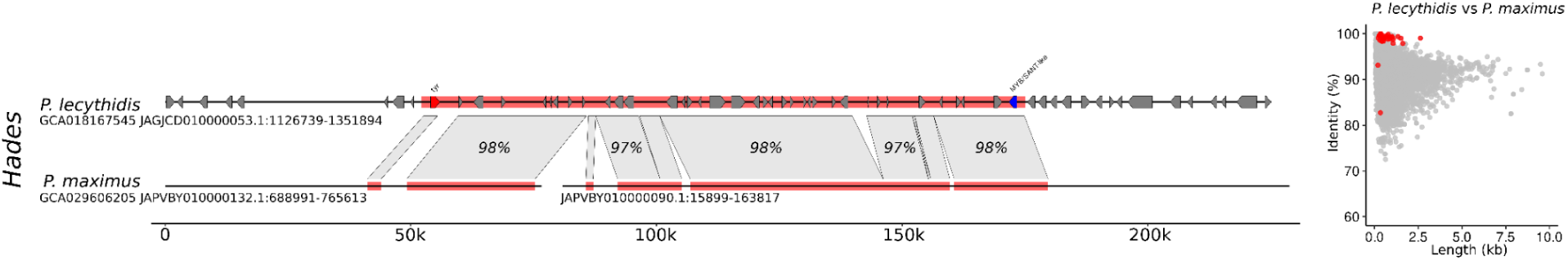
Synteny alignments (left) for the *Hades Starship* shown for species with evidence of its horizontal transfer. Predicted genes are depicted as arrows. *De novo* gene prediction was performed only for *Paecilomyces* isolates. BLASTall plots (right) showing nucleotide BLAST alignment length and identity using *P. lecythidis* protein-coding gene sequences as queries and whole genome sequence from *P. maximus* as a target. Predicted genes from *Hades* are shown in red.

**Figure S13:**
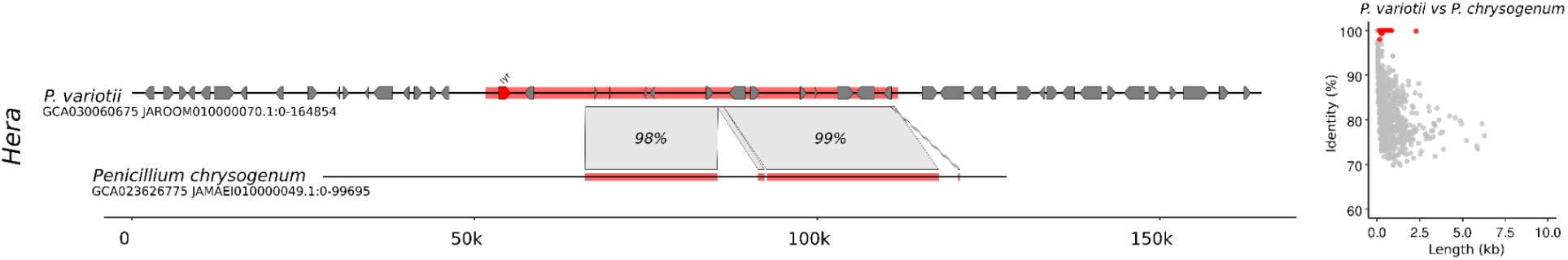
Synteny alignments (left) for the *Hera Starship* shown for species with evidence of its horizontal transfer. Predicted genes are depicted as arrows. *De novo* gene prediction was performed only for *Paecilomyces* isolates. BLASTall plots (right) showing nucleotide BLAST alignment length and identity using *P. variotii* protein-coding gene sequences as queries and whole genome sequence from *P. chrysogenum* as a target. Predicted genes from *Hera* are shown in red.

**Figure S14:**
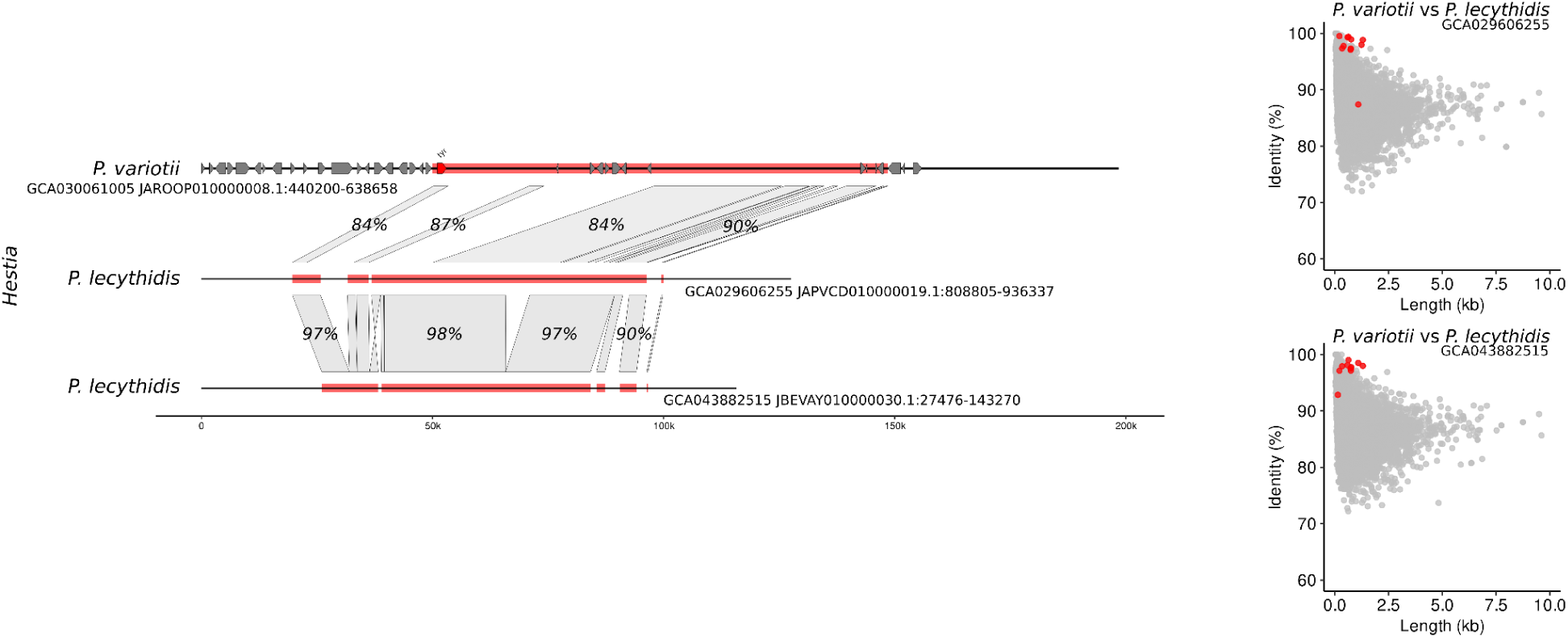
Synteny alignments (left) for the *Hestia Starship* shown for species with evidence of its horizontal transfer. Predicted genes are depicted as arrows. *De novo* gene prediction was performed only for *Paecilomyces* isolates. BLASTall plots (right) showing nucleotide BLAST alignment length and identity using *P. variotii* protein-coding gene sequences as queries and whole genome sequences from *P. lecythidis* as targets. Predicted genes from *Hestia* are shown in red.

**Figure S15:**
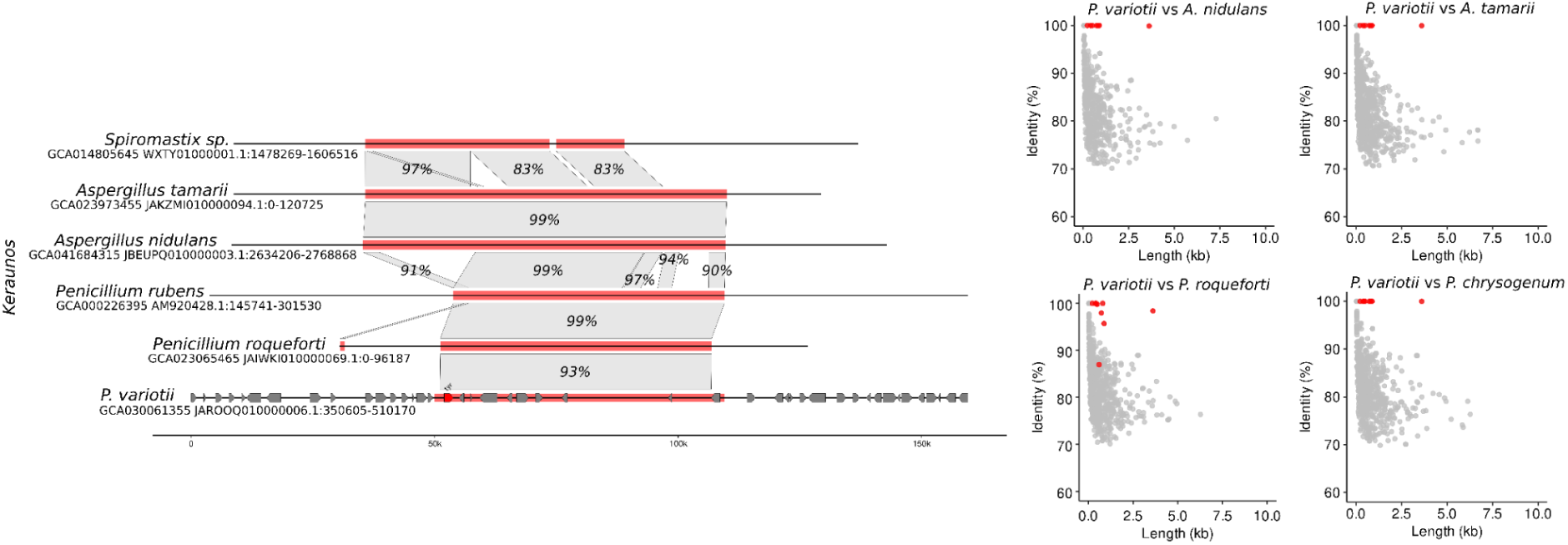
Synteny alignments (left) for the *Keraunos Starship* shown for species with evidence of its horizontal transfer. Predicted genes are depicted as arrows. *De novo* gene prediction was performed only for *Paecilomyces* isolates. BLASTall plots (right) showing nucleotide BLAST alignment length and identity using *P. variotii* protein-coding gene sequences as queries and whole genome sequences from the other named isolates as targets. Predicted genes from *Keraunos* are shown in red.

**Figure S16:**
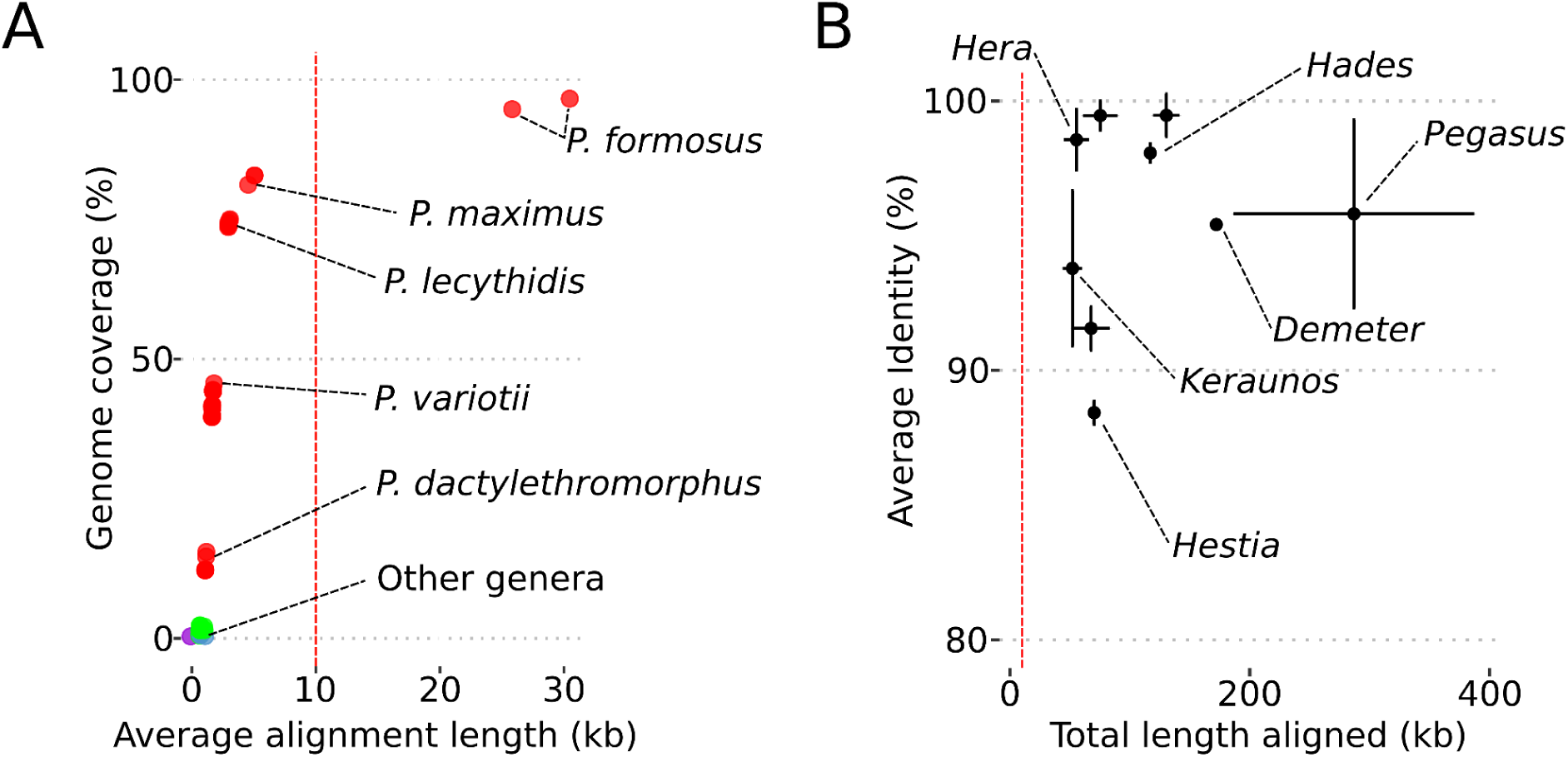
Summary statistics from alignments between whole genomes and *Starship* Haplotypes. **A)** Pairwise whole-genome alignments of a single focal *P. formosus* genome against all genomes within the *Paecilomyces* phylogeny showing how alignment coverage across the genome and average alignment length decreases with increasing phylogenetic distance. Red vertical line indicates a 10kb threshold that was used to filter out alignments in the HGT screen. **B)** Average identity and total length of alignments between *Starship* copies found across the different genomes predicted to have exchanged these 9 different *Starships* through HGT.

**Figure S17:**
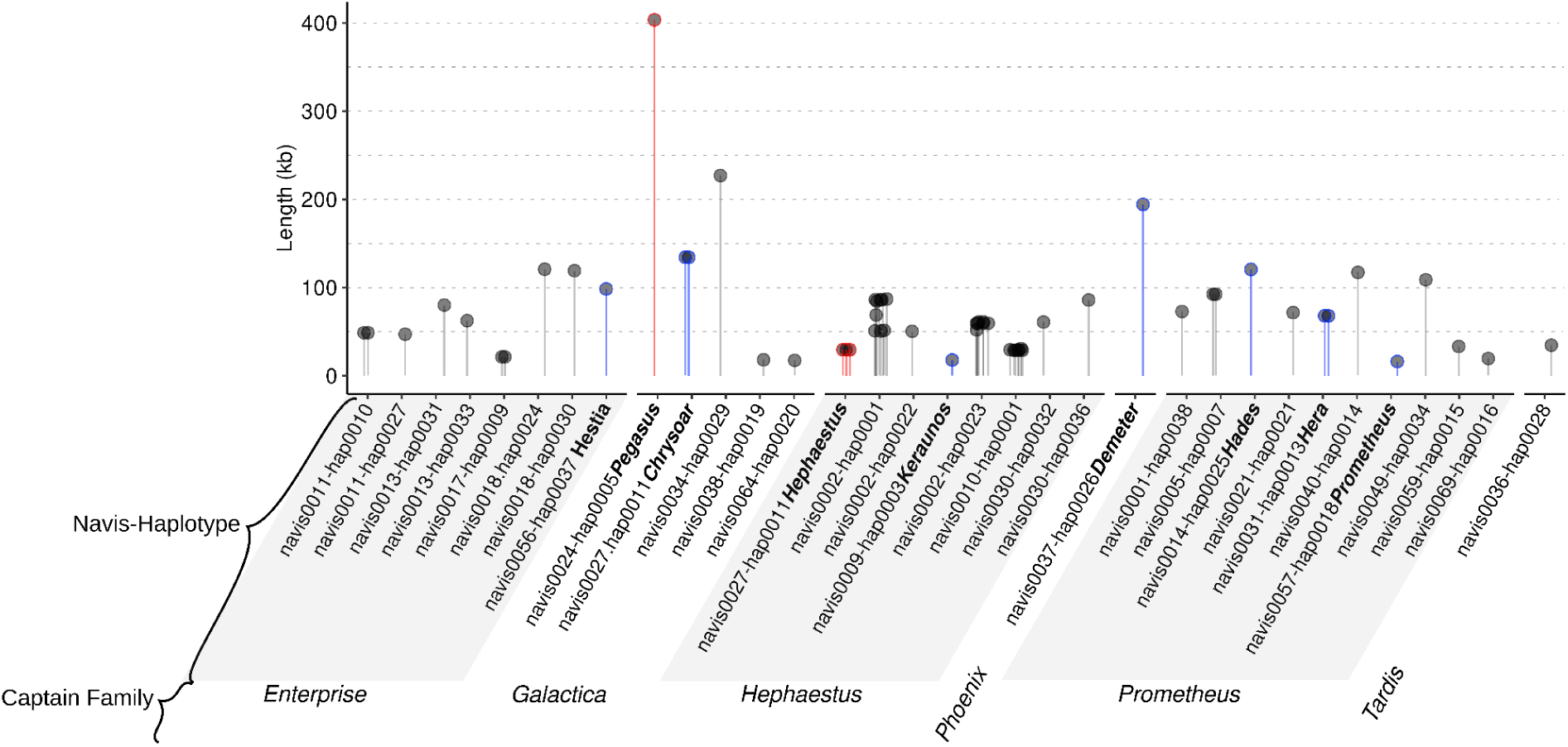
Length of 33 high-quality *Starships* identified by *Starfish* in the Paecilomyces. Both elements observed to have transferred experimentally are highlighted in red. All other *Starships* with evidence of HGT are highlighted in blue.

**Figure S18:**
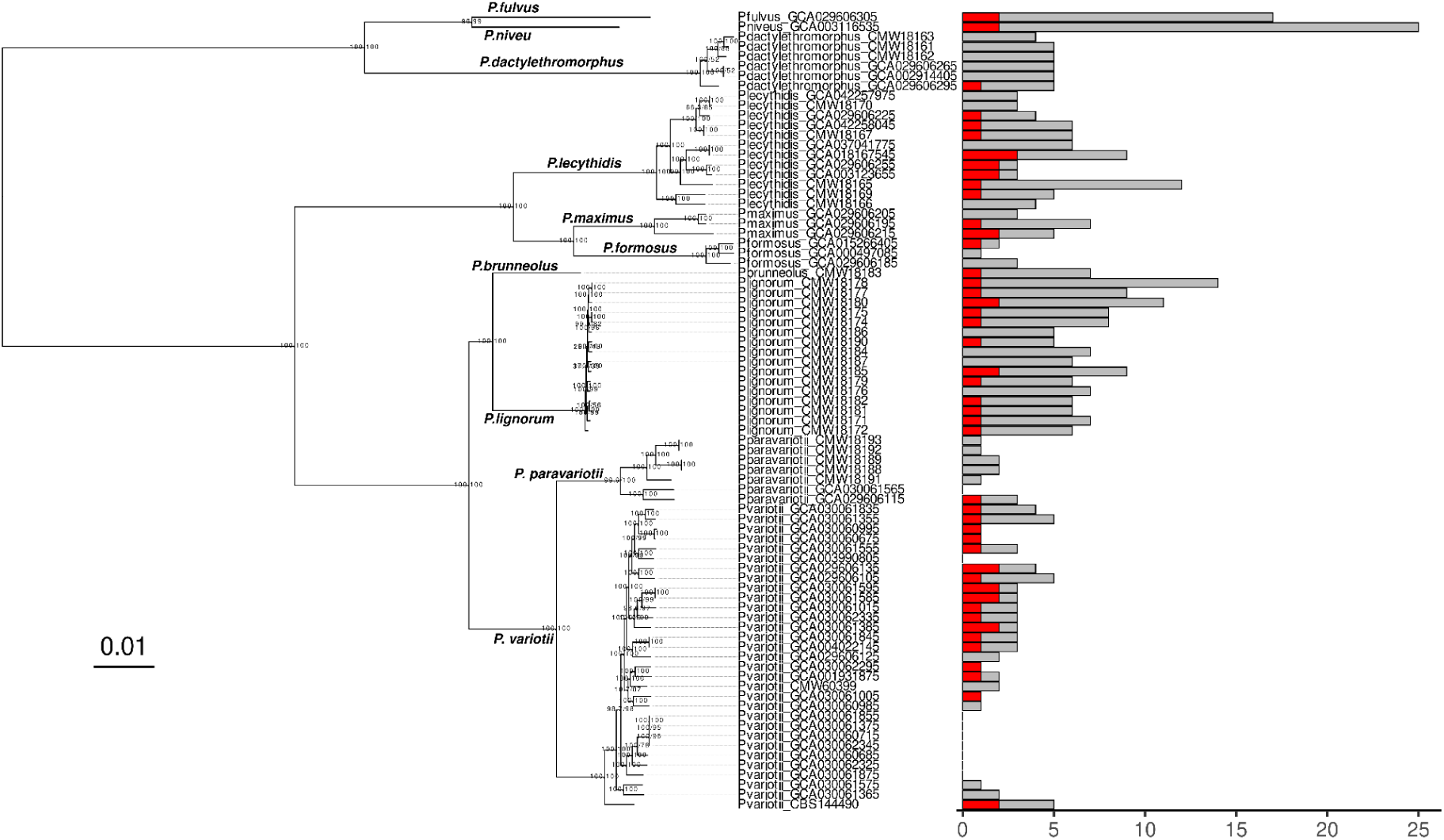
The total number of annotated captain tyrosine recombinases and the total number of full-length high-quality *Starships* per *Paecilomyces* genome, as predicted by starfish v1.1. The maximum likelihood phylogeny (left) was generated using conserved BUSCO orthologs from the 79 *Paecilomyces* genomes and midpoint rooted. Ultra-fast bootstrap and aLRT support percentages (out of 1000) are shown below each branch. The bar chart shows the total number of annotated captains (grey) and *Starships* (red) per genome.

## Notes

### Competing Interest Statement

The authors have declared no competing interest.

